# MiniCARbids: Minimalistic human binding domains specifically tailored to CAR T applications

**DOI:** 10.1101/2025.09.09.675083

**Authors:** Magdalena Teufl, Julia Mayer, Kerstin E. Holzer, Dominik Emminger, Elise Sylvander, Hayeon Baik, Fabian Schubert, Evangelia Maniaki, Ulrika Bader, Álvaro Muñoz-López, Joerg Mittelstaet, Markus Dobersberger, Michelle C. Buri, Leander Sützl, Benjamin Salzer, Eva M. Putz, Charlotte U. Zajc, Manfred Lehner, Michael W. Traxlmayr

**Affiliations:** Department of Chemistry, Institute of Biochemistry, BOKU University, Vienna, Austria; CD Laboratory for Next Generation CAR T Cells, Vienna, Austria; St. Anna Childreńs Cancer Research Institute, Vienna, Austria; Miltenyi Biotec B.V. & Co. KG, Bergisch Gladbach, Germany; Faculty of Life Sciences, Reutlingen University, Germany; Department of Food Science and Technology, Laboratory of Food Biotechnology, BOKU University, Vienna, Austria; Center for Physiology and Pharmacology, Medical University of Vienna, Vienna, Austria; Department of Transfusion Medicine and Cell Therapy, Medical University of Vienna, Vienna, Austria; St. Anna Children’s Hospital, Department of Pediatrics, Medical University of Vienna, Vienna, Austria

## Abstract

Traditionally, chimeric antigen receptor (CAR) T cells employ single-chain variable fragments (scFvs) as binding entities. While scFvs represent a convenient option due to their broad availability, they also come with drawbacks, in particular their tendency to cluster and their relatively large size. Moreover, most scFvs used in the CAR field are of non-human origin, potentially causing immunogenicity. Therefore, we established an engineering platform for minimalistic CAR binding domains (miniCARbids), which combine several critical advantages: (i) human origin, (ii) small size, (iii) efficient expression in T cells and (iv) single-domain architecture, among others. We demonstrate that miniCARbids can be engineered to recognize various antigens with antibody-like affinities, while being stable and aggregation-resistant. When miniCARbids are incorporated into CARs, they induce high anti-tumor potency in both adapter and conventional CAR formats. Remarkably, CD22-directed miniCARbid-based CARs showed similar or even more efficient tumor clearance in leukemia-bearing mice when compared with a CAR comprising the clinically tested m971-1xG_4_S scFv. Together, we introduce the miniCARbid engineering platform, enabling the generation of small, human antigen-binding domains with high potency in CAR T cells against virtually any target antigen.

## Introduction

Chimeric antigen receptor (CAR) T cells have revolutionized the treatment of hematologic malignancies including B cell acute lymphoblastic leukemia (B-ALL), B cell derived lymphomas and multiple myeloma.^1–4^ However, despite their great success, current CAR T cell therapies suffer from certain limitations such as resistance caused by antigen loss^1,3^ and limited efficacy due to T cell exhaustion,^5,6^ among others.

One central element determining CAR T cell efficacy and specificity is the CAR antigen binding domain commonly based on single-chain variable fragments (scFvs), which are usually used as the default option. However, while scFvs are convenient options that can readily be generated from existing monoclonal antibodies (mAbs), they also come with some disadvantages. As has been well-established in the antibody field, scFvs are known to form oligomeric structures due to domain swapping with neighboring molecules, resulting in the formation of diabodies, triabodies or tetrabodies.^7,8^ Importantly, this tendency of scFvs to cluster has also been observed on the surface of CAR T cells,^6,9^ which has been suggested to lead to antigen-independent tonic CAR signaling and – as a consequence – T cell exhaustion and dysfunction.^5,6^ This tendency to swap domains with neighboring scFvs is even more problematic in OR-gated CAR T cells expressing two scFvs in tandem.^10^ Apart from oligomerization tendencies, this two-domain structure also results in a considerable size of ∼250 amino acids, which is sometimes a limitation with respect to the packaging size of lenti-or retroviruses.^11–13^ Moreover, many currently used scFvs – and, in fact, all scFvs in CAR T products FDA-approved to date^1^ – are not of human origin, raising concerns about potential immunogenicity, which has indeed been demonstrated in several patients.^14,15^

To circumvent these limitations, alternative binder scaffolds including DARPins, nanobodies, Sso7d and monobodies have been tested in CAR T cells,^9,16–21^ demonstrating that CAR T cell function can also be achieved with non-antibody-based binding domains. While many of these studies have shown promising results with CARs based on non-scFv binding domains, none of these previously existing binder scaffolds had been designed specifically for CAR T cells and therefore their features are not ideal for this type of application. A phenomenon that is frequently observed with alternative binder scaffolds is the so-called stability-function trade-off, i.e. a considerable drop in stability during the engineering process, which in many cases leads to poor expression and/or aggregation,^20,22–25^ both of which are problematic in the context of CAR T cells.^5,6^ Moreover, the vast majority of these binder scaffolds are based on non-human proteins, potentially causing immunogenicity and rejection by the host immune system.^15,21^

Of note, any given stable protein can be engineered to turn into an antigen binding domain by using state-of-the-art protein engineering technologies. Therefore, in this study, we selected binding domains solely based on desired properties for applications in CAR T cells, including human origin, high expression levels in human CAR T cells and being based on a single protein domain to prevent clustering and excessive tonic signaling. Thus, these “minimalistic CAR binding domains” (miniCARbids) integrate all features required for efficient CAR function, while avoiding unnecessary payload.

To demonstrate the versatility of the miniCARbid platform, we have chosen three antigens of high relevance in the CAR T cell field: (i) CD22 (Siglec-2), which is a B cell lineage antigen commonly used for targeting of B cell-derived malignancies;^26,27^ (ii) CD276 (B7-H3), a checkpoint molecule overexpressed on a range of human tumors including neuroblastoma, sarcomas and brain tumors;^28^ and (iii) a peptide antigen for use in adapter CARs (AdCARs).^29,30^ For each of these antigens, we engineered stable and monomeric miniCARbids, with the majority reaching antibody-like affinities in the pM to nM range. Moreover, incorporation of these miniCARbids into 2^nd^ generation CARs resulted in potent CAR T cell activity for both conventional CAR, as well as AdCAR architectures. Remarkably, in addition to their small size saving vector payload and their human origin, miniCARbid-CARs showed similar or even more efficient elimination of leukemia in an *in vivo* model when compared with CARs based on the clinically tested m971 scFv, demonstrating the high potential of miniCARbids for use in CAR T cell therapies.

## Results

### Identification of binder scaffolds with ideal properties for CAR T cell applications

To establish a platform for the generation of binding domains specifically tailored to CAR T cell applications, we identified the following list of essential features (Fig. 1A): (i) human origin to reduce the risk of immunogenicity, (ii) small size to reduce vector payload, (iii) single-domain architecture to prevent domain mispairing as observed with scFvs,^5–9^ (iv) high stability to accommodate for the stability loss typically observed during the subsequent engineering process,^25^ (v) no aggregation, (vi) lack of cysteines to enable extra- and intracellular applications, (vii) lack of N-glycosylation motifs, (viii) availability of a surface exposed rigid β-sheet for binding site engineering, since this structural motif has been shown to yield binding domains with highly beneficial biochemical properties,^20,31–35^ (ix) intracellular origin (since therapeutically used human proteins sometimes induce antibody responses that cross-react with the endogenous protein,^36,37^ the binder scaffold should be derived from an intracellular protein, which is not accessible to such potential cross-reactive antibodies), (x) efficient expression on primary human T cells when fused to a CAR and (xi) no or only low tonic CAR signaling (Fig. 1A).

**Figure 1:**
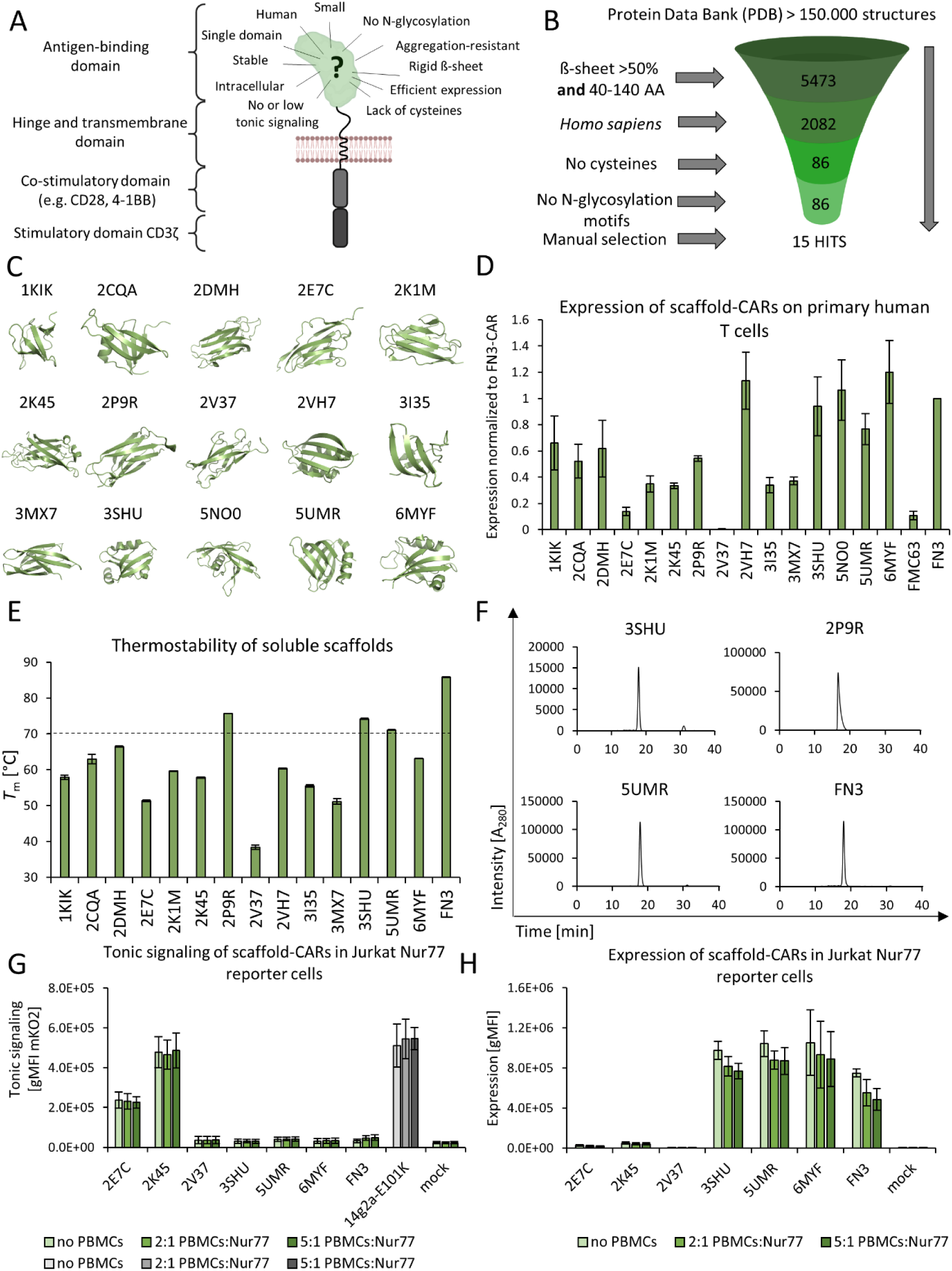
Selection process of novel binder scaffolds with ideal properties for CAR T applications. (A) Second-generation CAR molecule with a prospective antigen binding domain with the desired characteristics. (B) Selection process and criteria applied for the identification of novel potential scaffolds starting from the Protein Data Bank (PDB). (C) Structures of the 15 chosen potential scaffolds labeled according to their PDB-IDs. (D) Expression of 15 scaffold-CARs and FMC63-CAR and FN3-CAR as benchmarks on primary human T cells from 2 donors in a BBζ backbone (Q65K) (average ± SD, n=4, biological replicates). (E) 14 potential scaffolds and FN3 were expressed solubly and characterized regarding their thermostability using DSC; average ± SD of 3-4 independent measurements, technical replicates. (F) Aggregation properties were assessed using SEC-HPLC. Chromatograms from one representative experiment (n=3, technical replicates) are shown. (G) Tonic signaling of scaffold-CARs in a BBζ backbone (Q65K) in Jurkat Nur77 reporter cells was assessed based on the expression of mKusabira Orange (mKO2) in the absence or presence of PBMCs in 2-fold or 5-fold excess (average ± SD, n=3, biological replicates). A 14g2a-E101K-CAR was used as a positive control. (H) Expression levels of the scaffold-CARs shown in (G) (average ± SD, n=3, biological replicates). Expression was detected via anti-Flag-tag staining. Parts of this figure were created with BioRender.com.

To identify potential candidates fulfilling this extensive list of criteria, we screened all structures deposited in the Protein Data Bank (PDB). Applying the parameters defined in Fig. 1B yielded a list of 15 promising binder scaffold candidates (Fig. 1C; proteins labeled according to their PDB-IDs), which were subsequently fused to 4-1BB-based 2^nd^ generation (BBζ) CARs and tested for expression on primary human T cells. CARs containing the clinically used FMC63 scFv and the well-known alternative binder scaffold 10^th^ type III human fibronectin domain (FN3)^38,39^ within the same CAR backbone were included as references. While the expression rates varied greatly, most scaffold-CARs showed superior expression when compared with an identical CAR based on the FMC63 benchmark scFv (Fig. 1D and Suppl. Fig. 1A). Next, all candidates were expressed as soluble proteins and analyzed with respect to their thermal stabilities and aggregation tendencies (Fig. 1E, 1F and Suppl. Fig. 1B). Apart from the FN3 reference protein, only three proteins (2P9R, 3SHU and 5UMR) met the desired *T*_m_ of 70 °C. Since 2P9R showed peak tailing in size exclusion chromatography (SEC) analysis (Fig. 1F), presumably caused by a monomer-dimer equilibrium, we identified the intracellular proteins 3SHU (PDZ3 domain of the human tight junction protein ZO-1) and 5UMR (N-terminal domain of human FACT complex subunit SSRP1) as the most promising binder scaffold candidates. Further analysis of scaffold-CARs in Jurkat Nur77 reporter cells showed that – in contrast to the 14g2a-E101K scFv-based control CAR^40^ and the poorly expressed 2E7C and 2K45-based CARs – no or only very low tonic signaling was observed with 3SHU- and 5UMR-CARs (Fig. 1G), despite high expression levels (Fig. 1H).

Together, we identified the human proteins 3SHU and 5UMR as highly promising scaffolds for the engineering of minimalistic CAR binding domains (miniCARbids), integrating all desired properties defined above.

### Establishing randomly mutated miniCARbid libraries

To facilitate the generation of miniCARbids containing an engineered binding surface, a patch of surface exposed residues needed to be randomly mutated, followed by selection of antigen-specific miniCARbids from this highly diverse library. Since the introduction of mutations typically impairs protein folding and stability,^25^ it is of utmost importance to identify mutation-tolerant amino acid positions. For that purpose, we conducted structural, as well as phylogenetic analysis (Suppl. Fig. 2) and selected residues which are surface-exposed, non-conserved throughout evolution and which are not involved in stabilizing interactions with other residues within the domain.

Based on these parameters, we established NNK-randomized model libraries containing different sets of randomized positions (Fig. 2A) and analyzed them with respect to their full-length expression in the yeast display format, which has been shown to correlate with protein stability.^38,41,42^ Similar trends were observed after expression at 20 or 37 °C. For 3SHU, we chose library 3SHU_3 due to its efficient expression (Fig. 2B, Suppl. Fig. 3A) and favorable topology of its randomized positions, which form a contiguous binding pocket (Fig. 2A). For 5UMR, library 5UMR_c showed high level expression (Fig. 2B, Suppl. Fig. 3A), while library 5UMR_cp carries the advantage of an extended binding surface (Fig. 2A) that is still condensed onto a rather limited sequence stretch, thus limiting the number of potentially immunogenic peptides. Therefore, we chose to proceed with a library containing randomized positions of both 5UMR_c and 5UMR_cp, i.e. the positions of 5UMR_c and partially randomized positions Q28 and Q44 with a bias towards wild type glutamine.

**Figure 2:**
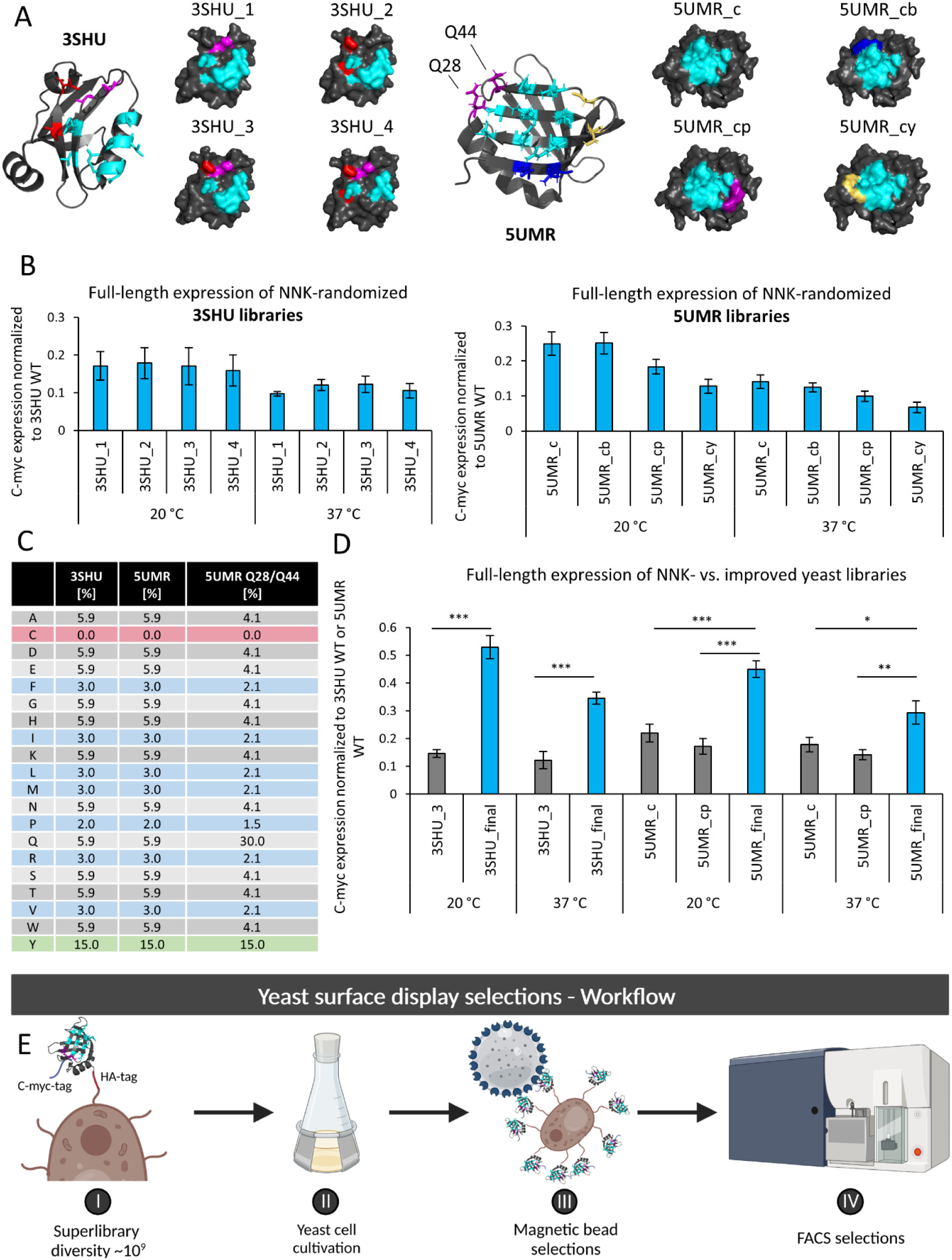
Establishment of miniCARbid libraries for binding site engineering. (A) Crystal structures of scaffolds 3SHU and 5UMR and four different library formats per scaffold with their mutated amino acids highlighted in color. (B) Full-length yeast surface expression of NNK-randomized 5UMR and 3SHU libraries normalized to the expression of 5UMR or 3SHU WT, respectively (induced at 20 and 37 °C) (average ± SD, n=3 or 4, biological replicates). (C) Amino acid frequencies chosen for the randomized positions in the optimized libraries, which differed for 5UMR positions Q28 and Q44. (D) Full-length expression of NNK-randomized vs. improved (final) yeast libraries normalized to the expression of 3SHU or 5UMR WT, induced at 20 and 37 °C (average ± SD, n=3, biological replicates). Statistical analysis was performed with a two-sided t-test for samples with equal variance (*p < 0.05, **p < 0.01, ***p < 0.001). (E) A typical workflow performed for yeast surface display selections starting from a superlibrary made up of scaffold libraries based on 3SHU and 5UMR with a total diversity of ∼10^9^. Parts of this figure were created with BioRender.com.

To further improve library quality, we used primers generated by trinucleotide synthesis, which allows for precise control of amino acid distributions within the randomized positions and exclusion of stop codons that are otherwise accidentally integrated in NNK-randomized libraries. We chose to increase the frequency of Tyr (15%) since this residue is known to be highly beneficial for antigen recognition,^43,44^ to reduce the percentage of Arg, Pro and hydrophobic amino acids and to exclude Cys residues (Fig. 2C). Furthermore, the 5UMR library contained 30% wild type residues at positions Q28 and Q44. Sequencing results of individual clones from these optimized libraries closely resembled the intended amino acid distributions (Suppl. Fig. 3B, 3C and 3D). Remarkably, this library optimization resulted in considerably improved performance of these advanced libraries termed 3SHU_final and 5UMR_final when compared with their NNK-randomized versions (Fig. 2D, blue vs. gray bars).

### Generation of stable and affine miniCARbids against multiple targets

Next, we mixed the two optimized libraries 3SHU_final and 5UMR_final to yield a superlibrary comprising two different binder scaffold topologies and a total diversity of ∼10^9^ randomly mutated variants. To efficiently select stable and affine miniCARbids, we employed the yeast surface display technology^45,46^ including magnetic bead selections and flow cytometric sorting (Fig. 2E) and three target antigens: CD22, CD276 (B7-H3) and a peptide antigen to be utilized in adapter CARs.

Selections against CD22 yielded multiple miniCARbids that recognize native CD22 on human leukemia cells with high affinities in the double digit pM to double digit nM range (Fig. 3A, 3B and Suppl. Fig. 4A). Furthermore, when expressed as soluble proteins, these CD22-miniCARbids were highly stable with *T*_m_ values between 51 and 71 °C (Fig. 3C). Three of the CD22-miniCARbids showed remarkably high *T*_m_ values comparable to the parental protein 5UMR, which is unusual for protein engineering campaigns that are typically associated with a significant loss in stability.^25^ Moreover, none of the miniCARbids showed detectable aggregation (Fig. 3D), further indicating the high quality of the optimized miniCARbid libraries. Finally, testing this set of miniCARbids for binding to CD22-negative (Jurkat) and CD22-positive (CD22^LOW^ NALM6 and CD22^HIGH^ Raji) cell lines demonstrated the high specificity of these engineered binding domains for CD22 (Fig. 3E).

**Figure 3:**
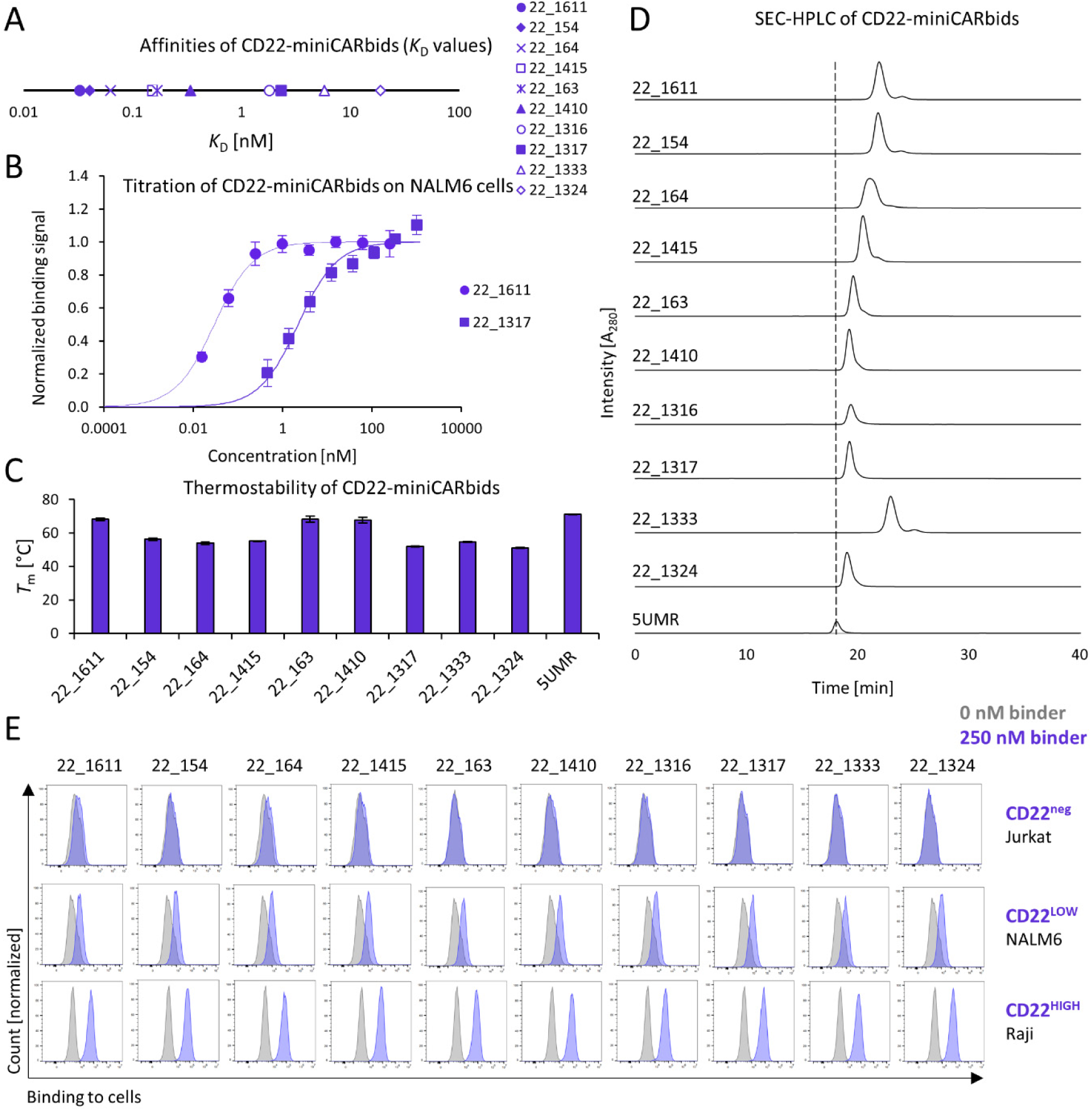
Engineered CD22-specific miniCARbids are affine, specific, stable and monomeric. (A) The *K*_D_ values of CD22-miniCARbids were determined by titrations of soluble CD22-miniCARbids on NALM6 cells. (B) A representative example of titrations of miniCARbids 22_1611 and 22_1317 on NALM6 cells is shown. The binding intensity was assessed via anti-His-tag staining by flow cytometry. Data were fitted with a 1:1 binding model (solid lines) for the calculation of the respective *K*_D_ values illustrated in (A) (average ± SD, n=3 or 4, biological replicates). (C) Thermostability of CD22-miniCARbids and their parental protein 5UMR was assessed using DSC (average ± SD of 3 independent measurements, technical replicates). (D) Aggregation properties of CD22-miniCARbids were assessed using SEC-HPLC. One representative analysis (n=3, technical replicates) of CD22-miniCARbids and their parental protein 5UMR is shown. (E) Binding specificity was assessed by incubating NALM6, Raji or Jurkat (CD22-negative) cells with 250 nM CD22-miniCARbid, followed by flow cytometric analysis (one of three biological replicates is shown).

As a second antigen, we chose CD276 which is frequently overexpressed on human cancers.^47,48^ After yeast display selections, we characterized eight enriched miniCARbids, three of which bound to CD276 on a panel of four human cancer cell lines (A-431, A-549, Caco-2, SK-BR-3; Fig. 4A and Suppl. Fig. 4B) known to express varying levels of the two CD276 isoforms 2Ig and 4Ig.^49^ Importantly, this interaction was specific, since no signal was obtained with CD276-negative Jurkat cells (Fig. 4B). Again, biochemical characterization of soluble miniCARbids revealed high stabilities (Suppl. Fig. 4C) and hardly any detectable aggregation (Fig. 4C and Suppl. Fig. 4E). Blocking experiments with well-established CD276-specific mAbs (enoblituzumab (MGA271) and omburtamab (8H9))^50^ showed different patterns for the three miniCARbids, suggesting that they bind to at least two different epitopes on CD276 (Suppl. Fig. 4D).

**Figure 4:**
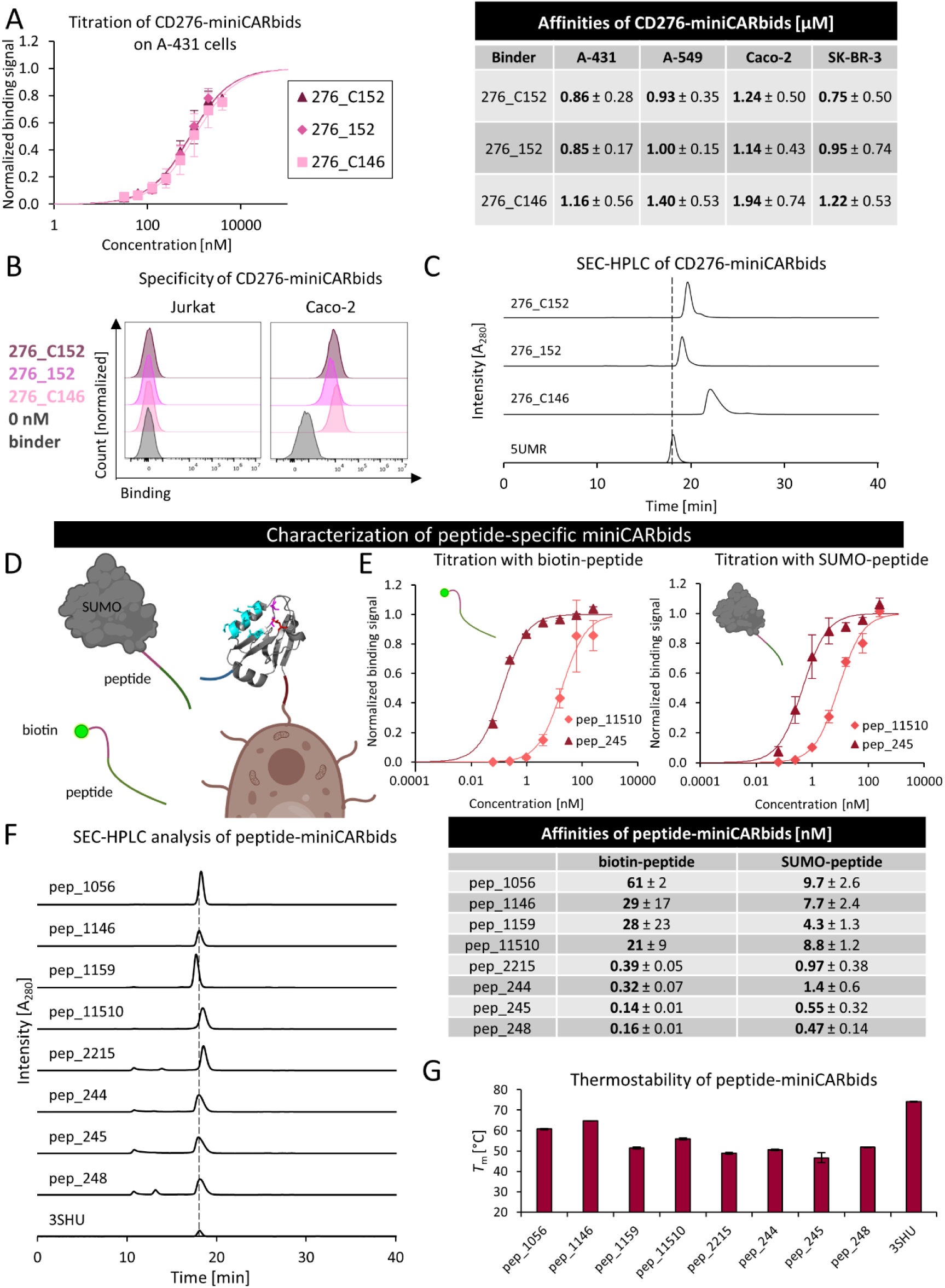
Engineered CD276- and peptide-specific miniCARbids show favorable biochemical properties. (A) *K*_D_ values and their respective titration curves of three CD276-miniCARbids (276_C152, 276_152 and 276_C146) on cell lines A-549 (14,140 ± 940 CD276 molecules/cell, average ± SD, n=3, biological replicates), A-431 (15,450 ± 4,110 CD276 molecules/cell, average ± SD, n=3, biological replicates), Caco-2 (45,200 ± 24,290 CD276 molecules/cell, average ± SD, n=3, biological replicates) and SK-BR-3 (2,040 ± 90 CD276 molecules/cell, average ± SD, n=3, biological replicates). The binding intensity was assessed via anti-His-tag staining by flow cytometry. Data were fitted with a 1:1 binding model (solid lines) for the calculation of the respective *K*_D_ values (average ± SD, n=3 or 4, biological replicates). (B) Binding specificity was assessed by incubating Caco-2 or Jurkat (CD276-negative) cells with 1000 nM CD276-miniCARbids, followed by flow cytometric analysis (n=3, biological replicates). (C) Analysis of aggregation properties of CD276-miniCARbids and their parental protein 5UMR by SEC-HPLC. One representative example (n=3, technical replicates) is shown. (D) Schematic representation of the titration of yeast-displayed peptide-miniCARbids with two antigens, SUMO-peptide and biotin-peptide. (E) Affinities (*K*_D_) and two representative titration curves of miniCARbids pep_11510 and pep_245 with antigens SUMO-peptide and biotin-peptide. The binding intensity was assessed via anti-His-tag staining or fluorescently labeled streptavidin by flow cytometry. Data were fitted with a 1:1 binding model (solid lines) for the calculation of the respective *K*_D_ values (average ± SD, n=3, biological replicates). (F) Analysis of aggregation properties of peptide-miniCARbids and their parental protein 3SHU by SEC-HPLC. One representative example of three independent measurements (technical replicates) is shown. (G) Thermostability of peptide-miniCARbids and their parental protein 3SHU was assessed using DSC (average ± SD of 3 independent measurements, technical replicates). Parts of this figure were created with BioRender.com.

To further challenge the versatility of the platform, we investigated whether miniCARbids can also be engineered for specific recognition of a short linear peptide antigen. Since these miniCARbids were intended to be used in adapter CARs (AdCARs), we chose a peptide antigen (the adapter) based on an oncogenic variant of fibroblast growth factor receptor 2 (FGFR2), which seems to escape immune surveillance in humans and still enables orthogonality due to the contained point mutations. Although the engineering of binder scaffolds against a short peptide comprising only several amino acids is considered quite challenging,^32^ we obtained miniCARbids that bound free peptide, as well as SUMO-peptide fusion protein with affinities in the low nM to high pM range (Fig. 4D and 4E and Suppl. Fig. 4F). Biochemical analysis showed that the miniCARbids were stable (Fig. 4G) and exhibited only little or no aggregation (Fig. 4F).

Together, these data demonstrate that miniCARbids can be engineered for specific recognition of very diverse antigens, including both proteins and linear peptides. All selection campaigns yielded stable and aggregation-resistant human binding domains of high affinity towards their target.

### High potency of CD22-specific miniCARbid-CAR T cells

Next, we investigated whether the miniCARbids are functional within conventional 2^nd^ generation CARs. For that purpose, we tested a set of ten CD22-specific miniCARbids, as well as three benchmark scFvs, two of which have been applied in CAR T cells in clinical trials (m971-1xG_4_S, m971-4xG_4_S).^26,27,51^ For proper comparison, all miniCARbids and scFvs were incorporated into the same CD28-based CAR backbone (28ζ, Fig. 5A). To directly assess CAR signaling, we used our recently established Jurkat Nur77 reporter cell line expressing monomeric mKusabira Orange2 (mKO2) under the control of the Nur77 promoter,^31^ which is indicative of early CAR and TCR activation.^52,53^ Among the ten miniCARbid-CARs, five showed expression levels comparable to those of the scFv-based CARs (Fig. 5B). Moreover, upon co-culture with CD22-positive target cells, all miniCARbid-CARs induced CAR activation to similar or even higher levels compared with scFv-based CARs (Fig. 5C). Interestingly, while all scFv-based CARs showed intermediate intensity of antigen-independent tonic signaling (Fig 5C, light grey bars), the set of ten miniCARbid-CARs yielded the full range of tonic signaling from virtually undetectable levels to strong antigen-independent activation (Fig. 5C, light blue bars). Of note, three miniCARbid-CARs (22_1410, 22_163 and 22_1611) showed efficient activation upon antigen engagement, despite negligible background activation in the absence of antigen (Fig. 5C, light blue vs. blue bars).

**Figure 5:**
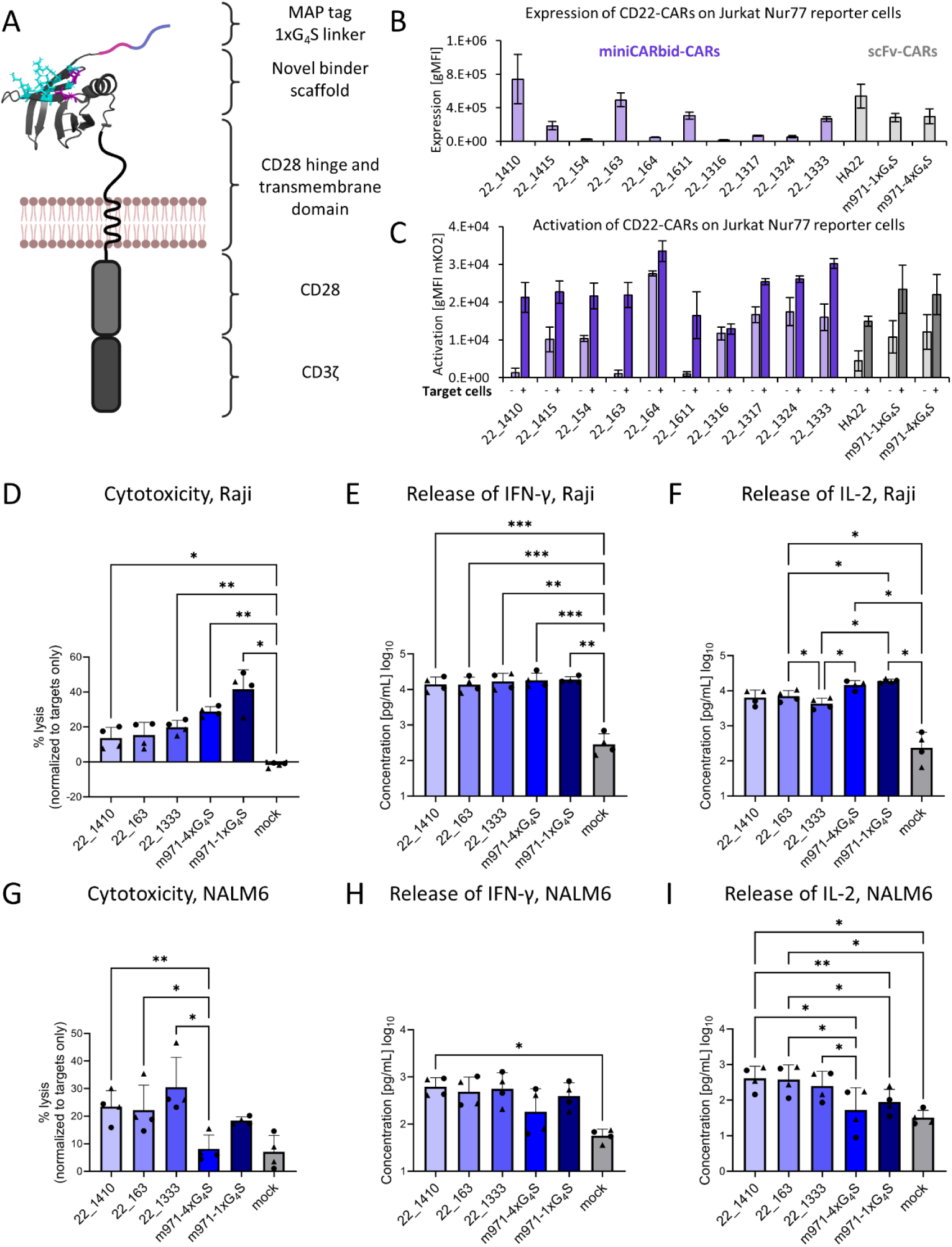
CD22-specific miniCARbid-CARs efficiently activate effector functions in human T cells. (A) CAR architecture used for the *in vitro* assessment of CAR activity. (B) Expression of CARs based on ten CD22-specific miniCARbids and scFvs HA22, m971-1xG_4_S and m971-4xG_4_S as benchmarks in Jurkat Nur77 reporter cells was assessed via anti-MAP-tag staining by flow cytometry (average ± SD, n=3, biological replicates). (C) Activation of CD22-specific CARs in Jurkat Nur77 reporter cells in the presence or absence of a 2-fold excess of NALM6 target cells was assessed via the expression of mKO2 by flow cytometry (average ± SD, n=3, biological replicates). (D) Cytotoxicity of CD22-specific CAR T cells and mock T cells (no CAR) against Raji cells (E:T 2:1, average ± SD, n=4, biological replicates). (E and F) Release of IFN-γ (E) and IL-2 (F) analyzed via ELISA. The cytokines were analyzed in the supernatants of co-cultures with Raji cells (E:T 2:1, average ± SD, n=4, biological replicates). (G) Cytotoxicity of CD22-specific CAR T cells and mock T cells (no CAR) against NALM6 cells (E:T 2:1, average ± SD, n=4, biological replicates). (H and I) Release of IFN-γ (H) and IL-2 (I) analyzed via ELISA. The cytokines were analyzed in the supernatants of co-cultures with NALM6 cells (E:T 2:1, average ± SD, n=4, biological replicates). Statistical analysis was performed using a repeated measure One-Way ANOVA with a Tukey post hoc test (*p < 0.05, **p < 0.01, ***p < 0.001). The statistical analysis for the cytokine concentration was performed using log-transformed values. Parts of this figure were created with BioRender.com.

To further confirm functionality of miniCARbid-CARs, we measured cytotoxicity and cytokine release in primary human T cells with the three most promising miniCARbid-CARs, as well as CARs based on the two clinically tested m971 scFvs. All CARs were efficiently expressed at similar levels (Suppl. Fig. 5G). When using Raji target cells, which express high levels of CD22 (∼25,000 molecules/cell, Suppl. Fig. 5H), all three miniCARbid-CAR Ts showed efficient cytotoxic activity (Fig. 5D and Suppl. Fig. 5D) and cytokine secretion. While there was a trend toward lower IL-2 release for miniCARbid-CAR Ts when compared with both m971-based CAR Ts, IFN-γ levels were similar (Fig. 5E and 5F and Suppl. Fig. 5E and 5F).

Since it is known that CD22 downregulation is a frequently observed escape mechanism in response to CD22-CAR Ts,^54^ highly sensitive CD22-CARs responding to low antigen levels are of critical importance. Therefore, we additionally assessed CAR T cell activity in response to NALM6 cells expressing low levels of CD22 (∼1,000 molecules/cell, Suppl. Fig. 5H) representative of expression levels of CD22 seen in ALL.^26,55^ Remarkably, we observed a trend toward more efficient lysis and cytokine release induced by the three miniCARbid-CARs compared with the m971 scFv-based CARs (Fig. 5G, 5H and 5I and Suppl. Fig. 5A, 5B and 5C).

Overall, CD22-specific miniCARbid-CARs trigger efficient target cell lysis and cytokine release. Importantly, our data also suggest that these CD22-specific miniCARbid-CARs are highly sensitive, enabling recognition and potent lysis of CD22^LOW^ tumor cells.

### CD276-directed miniCARbid-CAR T cells show efficient cytotoxicity and cytokine release

To test whether high CAR T cell functionality can also be obtained with miniCARbids directed against a different antigen, we tested three CD276-specific miniCARbids, as well as two clinically relevant benchmark scFvs (MGA271 and 376.96)^56,57^ in the same 28ζ CAR (Fig. 6A). Analysis of CAR activation in the Jurkat Nur77 reporter cell line demonstrated that one out of three tested miniCARbid-CARs (276_C152) showed expression comparable to the MGA271 scFv-based CAR (Fig. 6B). All miniCARbid-CARs triggered efficient antigen-dependent CAR activation, whilst inducing only little tonic signaling comparable to the MGA271 scFv-based CAR (Fig. 6C).

**Figure 6:**
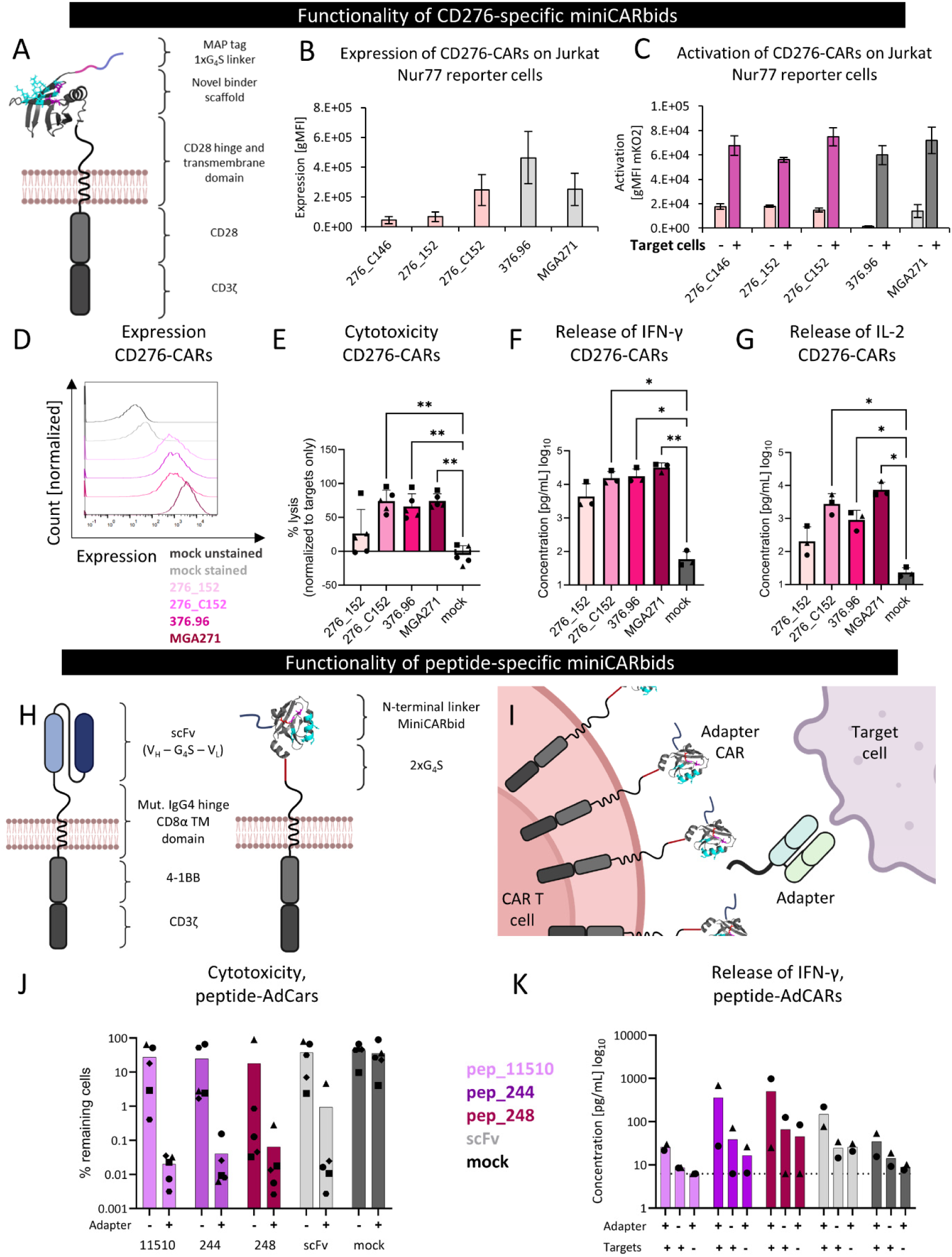
Efficient activation of CD276-specific miniCARbid-CARs and miniCARbid-based adapter CARs. (A) CAR architecture used for the *in vitro* assessment of CD276-specific miniCARbids. (B) Expression of CARs based on CD276-specific miniCARbids and scFvs MGA271 and 376.96 as benchmarks in Jurkat Nur77 reporter cells was assessed via anti-MAP-tag staining by flow cytometry (average ± SD, n=3, biological replicates). (C) Activation of CD276-specific CARs in Jurkat Nur77 reporter cells in the presence or absence of a 2-fold excess of A-549 target cells was assessed via the expression of mKO2 by flow cytometry (average ± SD, n=3, biological replicates). (D) Expression level of transduced CD276-specific CAR T cells was assessed via anti-MAP-tag staining by flow cytometry. (E) Cytotoxicity of CD276-specific CAR T cells and mock T cells (no CAR) against Caco-2 cells (E:T 5:1, 24 h, average ± SD, n=5, biological replicates). (F and G) Release of IFN-γ (F) and IL-2 (G) analyzed via ELISA. The cytokines were analyzed in the supernatants of co-cultures with Caco-2 cells (E:T 5:1, 24 h, average ± SD, n=3, biological replicates). (H) CAR architecture used for the functional characterization of peptide-specific miniCARbids in a CAR format. (I) MiniCARbids as well as an scFv were tested in an AdCAR format with a soluble peptide-adapter molecule. CAR activity is anticipated only upon the addition of the adapter protein. (J) Cytotoxicity of peptide-specific CAR T cells and mock T cells (no CAR) against OCl-AML2 cells (E:T 2:1, 4 d, average, n=5, biological replicates). Three miniCARbids (pep_11510, pep_244, pep_248; pink, purple and red) were tested and compared with a benchmark scFv (light grey). (K) Release of IFN-γ analyzed via MACSPlex Cytokine Kit. The cytokines were analyzed in the supernatant 24 h after the initiation of co-culture with OCl-AML2 cells (E:T 2:1, average, n=2, biological replicates). The dotted line represents the detection limit. Parts of this figure were created with BioRender.com.

Next, the two most promising CD276-specific miniCARbid-CARs were assessed for cytotoxicity and cytokine release in primary human T cells. 6 and 24 h co-culture assays were performed with a panel of three different CD276-expressing human cancer cell lines (A-431, A-549 and Caco-2) at effector:target (E:T) ratios of 5:1 and 10:1. While miniCARbid-CAR 276_152 induced moderate lysis, as well as IFN-γ and IL-2 secretion upon co-culture with all target cell lines, CAR T cells based on 276_C152 showed high potency comparable to those achieved with the two scFv-based control CARs (Fig. 6D-G and Suppl. Fig. 6).

Overall, these data demonstrate that CD276-directed miniCARbid-CAR T cells are highly functional and comparable to benchmark scFv-based CAR T cells, as demonstrated by efficient CAR T cell signaling, cytotoxic activity and cytokine release.

### Adapter CAR (AdCAR) T cells based on miniCARbids prove to be functional

To further demonstrate the versatility of miniCARbids in different CAR T applications, peptide-directed miniCARbids were used to design BBζ AdCARs, where the miniCARbids served as recognition domains for the soluble adapter protein containing the mutant FGFR2 peptide, as well as a tumor-specific Fab (Fig. 6H and 6I). Primary human AdCAR T cells were co-cultured with the acute myeloid leukemia (AML) cell line OCl-AML2 in the presence or absence of CD33-specific soluble adapter protein and assessed for cytotoxicity, cytokine release and expression of activation markers (CD69 and CD137). Three peptide-directed miniCARbid-AdCARs were tested together with an scFv-based AdCAR specific for the same peptide, all of which showed target cell lysis upon addition of adapter protein that was comparable to that achieved with the scFv-based AdCAR (Fig. 6J). Adapter-dependent activation was further confirmed by detection of activation markers CD69 and CD137 (Suppl. Fig. 7A and 7B). Moreover, analysis of cytokine secretion demonstrated that AdCAR activation was dependent on the presence of both the soluble adapter and CD33-positive target cells (Fig. 6K and Suppl. Fig. 7C and 7D). Together, these data demonstrate that miniCARbids can also be engineered for specific recognition of a linear peptide, thus enabling the generation of miniCARbid-based AdCAR T cells.

**Figure 7:**
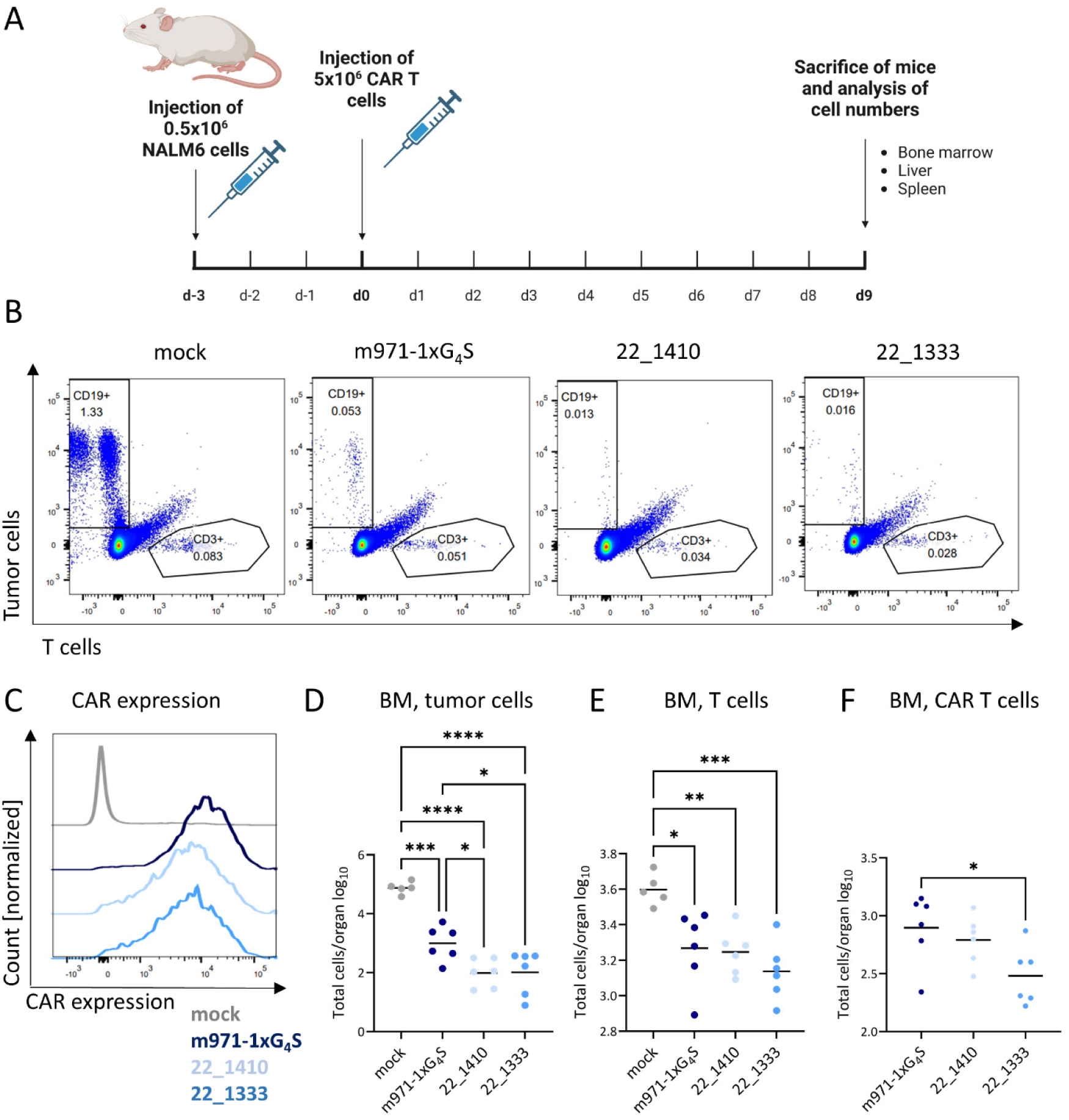
Pronounced anti-tumor potency of CD22-specific miniCARbid-CAR T cells *in vivo*. (A) Experimental design: Mice were injected with 0.5×10^6^ NALM6 cells, followed by the injection of 5×10^6^ CAR T cells three days later. On day nine post CAR T injection, mice were sacrificed and analyzed regarding the number of tumor, T and CAR T cells. (B) Flow cytometric analysis of bone marrow (BM) single-cell suspensions for tumor and T cell number; one representative sample for mock T cells, m971-1xG_4_S-CAR T cells and miniCARbid-CAR T cells 22_1333 and 22_1410 is shown. (C) CAR expression levels of injected CAR T cell suspensions as assessed via anti-MAP-tag staining. (D) Analysis of the number of tumor cells (CD19+ GFP+) of five (mock T cells) or six animals (CAR expressing T cells) in the bone marrow. Depicted is the log-transformed number of total cells in the bone marrow. (E and F) Analysis of the number of T cells (CD3+) (E) and of CAR T cells (MAP+) (F) of five (mock T cells) or six animals (CAR expressing T cells) in the bone marrow. Depicted is the log-transformed number of total cells in the bone marrow. Statistical analysis was performed using a One-Way ANOVA of log-transformed values with a Tukey post hoc test (*p < 0.05, **p < 0.01, ***p < 0.001, ****p < 0.0001). Parts of this figure were created with BioRender.com.

### High *in vivo* anti-tumor potency of CD22-specific miniCARbid-CARs

Finally, to assess the functionality of miniCARbid-CARs in a 28ζ CAR backbone *in vivo*, we compared the two most promising CD22-specific miniCARbid-CARs (22_1410 and 22_1333) to a CAR based on the m971 scFv. We chose the m971 scFv version with the shorter linker (1xG_4_S) as our benchmark, since it was shown to be more potent in clinical CAR T trials compared with the same scFv containing a longer linker (4xG_4_S),^27^ a trend that was further supported by our *in vitro* cytotoxicity assays (Fig. 5D and 5G). To mimic CD22^LOW^ tumors, we used a NALM6 model, in which 0.5×10^6^ NALM6 cells were injected three days prior to the administration of 5×10^6^ CAR T cells (CAR expression shown in Fig. 7C), followed by analysis of tumor and CAR T cell numbers in the bone marrow, liver and spleen at day nine post CAR T cell injection (Fig. 7A).

Importantly, both miniCARbid-CAR constructs induced potent killing of NALM6 tumor cells in the bone marrow (Fig. 7B, 7D and Suppl. Fig. 8A). Similarly, in the liver, where the tumor cell number was much lower in general, miniCARbid-CAR T cells also reduced the tumor burden (Suppl. Fig. 8B, only statistically significant for 22_1410). As was observed in previous studies,^9^ we did not detect NALM6 cells in the spleen (not shown). Remarkably, in the bone marrow, which showed the highest infiltration of NALM6 tumor cells, both miniCARbid-CAR T cells enabled significantly stronger eradication of tumor cells compared to the m971-scFv-based CAR Ts (Fig. 7B, 7D and Suppl. Fig. 8A), further demonstrating their high anti-tumor potency. Analysis of total, as well as CAR-positive T cell numbers in the bone marrow and liver showed comparable or slightly lower cell counts for the miniCARbid-CAR T cells when compared to the m971-CAR (Fig. 7E and 7F and Suppl. Fig. 8C and 8D).

Summing up, this *in vivo* experiment further confirmed the high efficacy of miniCARbid-CARs, which showed similar or even higher anti-tumor potency in leukemia-bearing mice when compared with a CAR based on the clinically tested m971-1xG_4_S scFv.

## Discussion

In this study, we established a platform for the generation of miniCARbids, i.e. small, stable and well-expressed human binding domains. Moreover, due to their single-domain architecture, the risk of clustering as frequently observed with scFvs is strongly reduced. Thus, while there is extensive experience with scFv-based CARs due to their broad availability, we present miniCARbids as an alternative platform that is specifically tailored to applications in CAR T cells.

We successfully engineered miniCARbids against three antigens, including two different proteins and a linear peptide. In all cases, we obtained stable miniCARbids with affinities down to the pM range, which is comparable to those typically obtained with antibodies. Thus, our data suggest that miniCARbids can be engineered for specific recognition of virtually any given antigen.

When engineering proteins to obtain new functionalities such as antigen binding, a loss of stability is commonly observed due to the insertion of mutations.^22,23,25,58^ Of note, some of the miniCARbids generated in this study only showed very minor loss in stability (difference in *T*_m_ of <3 °C compared to parental protein). Moreover, the vast majority of our engineered miniCARbids did not show any detectable aggregation, which is remarkable, as protein engineering for antigen recognition is typically associated with increased tendencies to aggregate.^20,22,58^ These highly favorable biochemical properties of the obtained miniCARbids, i.e. their high stability and low tendency to aggregate, prove the high quality of the library and mutational tolerance of the parental protein scaffolds. Moreover, these beneficial features may also at least partially explain the observed high performance of miniCARbids in CAR T cells. In this regard, it is worth noting that the miniCARbids obtained from yeast display selections were directly integrated into standard 2^nd^ generation CARs without any optimization of the CAR architecture (e.g. hinge length, signal peptide, etc.) or the miniCARbids themselves. Despite this omission of optimization efforts, the resulting miniCARbid-CARs were highly functional, comparable or even superior to well-established and extensively optimized scFv-benchmark CARs.

In fact, CD22-specific miniCARbid-CARs showed high sensitivity toward CD22^LOW^ NALM6 target cells, which resulted in superior anti-tumor activity in the bone marrow of mice *in vivo* when compared with a CAR based on the clinically tested m971-scFv (Fig. 7B and D). This finding is also of high clinical relevance, because CD22 downregulation has been shown to be an escape mechanism in leukemia and lymphoma patients upon treatment with CD22-directed CAR T products,^54,59^ suggesting that the high sensitivity of CD22-directed miniCARbids-CARs may impede tumor escape. In addition to this high potency and sensitivity, miniCARbids also come with the advantage of their human origin and smaller size (0.3 kbp vs. ∼0.75 kbp for a typical scFv), which considerably decreases the required vector payload that is generally known to significantly impact virus titer and infectivity.^11,12^ This reduced size is mostly attributed to the single-domain architecture of miniCARbids, which also prevents domain swapping and mispairing – a benefit that is particularly advantageous when expressed in a tandem-CAR format, where mispairing can be even more problematic due to the presence of two scFvs.^5,10^

Of note, due to their highly favorable properties, we anticipate that miniCARbids can also be utilized for a range of other applications, including bispecific T cell engagers (BiTEs), Fc fusions or drug-dependent switches, to name a few. For example, also in BiTEs, which are often based on two scFvs, the use of miniCARbids may prevent unintended domain swapping.

Together, the miniCARbids introduced in this study combine several critical features for applications in CAR T cells. First of all, miniCARbids are based on small protein domains of human origin. We demonstrate that these minimalistic single-domain proteins can be engineered to bind to diverse types of antigens with antibody-like affinities, while maintaining a stable and aggregation-resistant structure. In addition, even without optimization of the CAR architecture, miniCARbid-based CARs proved to be highly potent *in vitro* and *in vivo*, suggesting that this engineering platform will be a rich resource for small, human binding domains optimized for CAR T applications.

## Acknowledgements

This work was supported by the Austrian Science Fund (FWF Projects W1224 – Doctoral Program on Biomolecular Technology of Proteins – BioToP, 10.55776/P34832, ESP 465-B and EFP 45, Devising Advanced TCR-T cells to eradicate OsteoSarcoma, DART2OS) and by the Federal Ministry for Digital and Economic Affairs of Austria and the National Foundation for Research, Technology and Development of Austria to the Christian Doppler Research Association (Christian Doppler Laboratory for Next Generation CAR T Cells) and by private donations to the St. Anna Childreńs Cancer Research Institute (Vienna, Austria). E. S. and and M.C.B. are recipients of DOC Fellowships of the Austrian Academy of Sciences at the St. Anna Childreńs Cancer Research Institute (#26323 and #25905). The SH800S cell sorter, the CytoFLEX and the PEAQ-DSC Automated equipment was kindly provided by the EQ-BOKU VIBT GmbH and the project was supported by the BOKU Core Facility Biomolecular & Cellular Analysis.

## Author contributions

M.T. managed the project, conducted experiments, performed data and statistical analysis, prepared the figures and manuscript. J.M., K.E.H. and D.E. conducted experiments and performed data analysis. E.S. and H.B. conducted experiments and mouse work. F.S. conducted experiments and performed data analysis. E.M. and U.B. conducted experiments and performed data analysis. A.M.-L. and J.M. contributed to data interpretation. M.D. supported with data and statistical analysis. M.C.B. conducted mouse work. L.S. conducted experiments and performed data analysis. B.S., E.M.P. and C.U.Z. contributed to project conceptualization and data interpretation. M.L. contributed to the study idea and data interpretation. M.W.T. conceived the study idea, supervised the project, contributed to data interpretation and prepared the manuscript. All authors revised the manuscript.

## Data availability

All data are available upon request.

## Materials and Methods

### Expression and purification of soluble scaffolds and miniCARbids

All soluble scaffolds and miniCARbids were expressed in a pET-21a(+) vector (Novagen) with an N-terminal MGGGSGGSGG-linker, a C-terminal 2x(G_4_S)-G linker followed by a hexahistidine (6xHis) tag. Briefly, *E. coli* Tuner cells were transformed with sequence-verified plasmids and grown overnight in lysogeny broth (LB) with 100 µg/mL ampicillin at 37 °C, 180 rpm. Next, the culture was diluted to an OD_600_ of 0.1 – 0.2 in terrific broth and at OD_600_ ∼ 0.8-1 protein expression was induced with 1 mM isopropyl-beta-D-thiogalactopyranoside (IPTG) and the culture was shaken at 20 °C overnight. After centrifugation (5,000 g, 20 min, 4 °C), cells were resuspended in sonication buffer (50 mM sodium phosphate, 300 mM NaCl, 3% glycerol, 1% Triton-X 100, pH 8) and sonicated on ice, followed by centrifugation (20,000 g, 30 min, 4 °C) to separate the soluble proteins from the rigid cell matter.

Next, the proteins were purified by using TALON metal affinity resin (Takara Bio). Supernatants from crude cell lysates with 10 mM imidazole were applied to the washed and equilibrated TALON matrix twice. After several washing steps with equilibration buffer (50 mM sodium phosphate, 300 mM NaCl, pH 8) containing increasing amounts of imidazole (5 mM, 15 mM), the proteins were eluted with equilibration buffer containing 250 mM imidazole.

Buffer exchange to PBS was either performed using Amicon filter tubes (Merck Millipore) or by dialysis using SnakeSkin Dialysis Tubing (Thermo Fisher Scientific) at 4 °C. Protein concentration was determined using A_280_ and protein aliquots were frozen at −80 °C.

### Size exclusion chromatography (SEC) – HPLC

The aggregation behavior of soluble proteins was measured by SEC-HPLC (Shimadzu prominence LC20), equipped with a diode array detector (SPD-M20A, Shimadzu), by using a Superdex 75 10/300 column and PBS (additionally containing 200 mM NaCl) at a flow rate of 0.75 mL/min. 50 µg protein sample were loaded, unless stated otherwise.

### Differential scanning calorimetry (DSC)

DSC was performed using the automated MicroCal PEAQ-DSC (Malvern Panalytical). Proteins were measured at either 50 µM (5UMR, wildtype scaffolds) or 100 µM (3SHU) in PBS. Samples were heated from 20-100 or 120 °C at a heating rate of 60 °C/h. Data analysis was performed with the MicroCal PEAQ-DSC Software (Malvern Panalytical) performing a buffer baseline subtraction followed by normalization for protein concentration and fitting to a non-two state unfolding model.

### Cell culture

Buffy coats from de-identified healthy donors were purchased from the Austrian Red Cross. Primary human T cells were isolated by negative selection using the RosetteSep Human T Cell Enrichment Cocktail (STEMCELL Technologies) and cryopreserved in RPMI-1640 GlutaMAX medium supplemented with 20% (v/v) fetal calf serum (FCS, Sigma) and 10% DMSO. Upon thawing, T cells were washed with RPMI-1640 GlutaMAX medium and immediately activated using Human T-Activator CD3/CD28 dynabeads (Thermo Fisher Scientfic). T cells were counted every second day and were kept between a density of 0.3-1×10^6^ cells/mL. They were expanded for at least 7-10 days in RPMI-1640 GlutaMAX supplemented with 10% FCS (v/v), 100 U/mL penicillin, 100 µg/mL streptomycin and 200 U/mL of IL-2 (PreproTech).

Jurkat, Jurkat Nur77 reporter cells,^31^ NALM6, NALM6 GFP/luciferase, A-431, A-431 GFP/luciferase, A-549, A-549 GFP/luciferase, SK-BR-3, Raji and Raji GFP/luciferase cells were cultivated in RPMI-1640 GlutaMAX™ supplemented with 10% (v/v) FCS, 100 U/mL penicillin and 100 µg/mL streptomycin. Suspension cells were grown at densities between 0.3-1.5×10^6^/mL at 37 °C (5% CO_2_, 97% humidity. Caco-2 and Caco-2 GFP/luciferase cells were cultivated in DMEM supplemented with 20% (v/v) FCS, 100 U/mL penicillin and 100 µg/mL streptomycin. NALM6 cells, Raji and Jurkat cells were kind gifts from Dr. Sabine Strehl and Dr. Michael Dworzak, respectively; CCRI, Vienna, Austria. Caco-2 cells were a kind gift from Dr. Johannes Grillari, BOKU University, Vienna, Austria. A-549 (ATCC CCL-185), A-431 (ATCC CRL-1555) and SK-BR-3 (ATCC HTB-30) were obtained from ATCC. NALM6 GFP/luciferase, Raji GFP/luciferase, A-431 GFP/luciferase, A-549 GFP/luciferase and Caco-2 GFP/luciferase were established in-house.

### Expression of scaffold-CARs on primary human T cells

DNA encoding a T7 RNA polymerase promoter site, Kozak sequence and the transgene was PCR amplified and subsequently used as a template for *in vitro* transcription using the mMESSAGE mMACHINE T7 Ultra Kit (Thermo Fisher Scientific), followed by subsequent mRNA purification using the RNeasy kit (Qiagen). Purified mRNA was stored at −80 °C. 2×10^6^ primary human T cells were electroporated with 5 µg of scaffold-CAR mRNA and 1 µg of mRNA encoding for GFP using the Gene Pulser Xcell Electroporation system (Bio-Rad). Beforehand, cells were washed with RPMI-1640 without FCS, RPMI-1640 without phenol red and Opti-MEM (300 – 500 g, 5 min, RT) and the cell density was adjusted to 2×10^6^ cells/100 µL Opti-MEM. Electroporation was performed using the square wave protocol (single pulse, 500 V, 4 mm electroporation cuvettes, 5 ms pulse length) and cells were rescued directly after electroporation in 2 mL of pre-warmed full-growth medium (RPMI-1640 GlutaMAX, 10% FCS, 100 U/mL penicillin, 100 µg/mL streptomycin and 200 U/mL IL-2). After 16-18 hours of incubation 100,000 cells were used for flow cytometric analysis. After a washing step with 1 mL ice-cold staining buffer (PBS, 0.2% human albumin, 0.02% sodium azide) cells were blocked with 10% human serum for 10 min at 4 °C. 1.2 µg/mL anti-FLAG-PE antibody (BioLegend, clone L5) was added and cells were incubated for 25 min (in the dark, 4 °C) and subsequently washed twice with 1 mL ice-cold staining buffer. Samples were analyzed with a LSR Fortessa instrument (BD Biosciences).

### Tonic signaling using a Jurkat Nur77 reporter cell line

Preparation of mRNA and electroporation were performed as described above, using 2×10^6^ Jurkat Nur77 reporter cells and 5 µg mRNA and a square wave protocol (single pulse, 500 V, 4 mm, 3 ms pulse length). Cells were rescued in full-growth medium without IL-2. After 16-20 hours of incubation 50,000 Jurkat Nur77 reporter cells were co-cultured with 100,000 or 250,000 (1:2 or 1:5 E:T ratio) PBMCs (200 µL total volume) for 4 hours at 37 °C in 96-well U-bottom plates. After the incubation time, cells were centrifuged (300-500 g, 5 min, 4°C), washed with 200 µL ice-cold PBSA (PBS, 0.1% BSA) and stained with 1.2 µg/mL anti-FLAG-APC (BioLegend, clone L5) for 20 min (4 °C, shaking, in the dark). Jurkat Nur77 reporter cells expressing the 14g2a-E101-CAR were blocked with 100 µg/mL MOPC (IgG1, Kappa from murine myeloma, Sigma-Aldrich) for 10 min at 4 °C, followed by staining with 10 µg/mL of anti-IgG Fc-AF647 (Southern Biotech, clone JDC-10) for 20 min (4 °C, shaking, in the dark). After two washing steps, the samples were analyzed with a Cytoflex S instrument (Beckman Coulter).

### Phlyogenetic analysis of 3SHU and 5UMR to predict conserved residues

For 3SHU, covering a large sequence diversity without oversampling the dense local sequence space was achieved by using the full-length *H. sapiens* Tight junction protein ZO-1 (seq. ID: Q07157) as query for a search on the representative sequence database UniRef90. The search was conducted using the sequence BLAST option of the Enzyme Function Initiative-Enzyme Similarity Tool^60^ with default settings and the resulting network was finalized at an alignment score of 1×10^−50^. All sequences clustering with Q07157 and showing a sequence length between 30 and 2000 were further aligned by MAFFT v7.402^61^ using the FFT-NS-2 method. The alignment was trimmed for positions with >90% gaps using trimAl v1.2^62^ and manually trimmed to fit the length of the domain 3SHU. After removing sequence redundancy of >90%, but keeping 3SHU in the selection, conserved residues in the alignment were visualized using WebLogo 3.^63^

Investigation of the sequence space of 5UMR was conducted by running a PSI-BLAST search of the full-length *H. sapiens* FACT complex subunit SSRP1 (seq ID: NP_003137.1) and its homolog from *S. cerevisiae* (seq. ID: NP_013642.1) respectively, using a PSI-BLAST threshold of 9E-50 and three iterations. Resulting unique sequences from both searches combined were aligned by MAFFT v7.402^61^ using the FFT-NS-2 method and cut after the highly conserved ‘GWNWG’-motif to fit the length of the domain 5UMR. Sequences with a length <80, sequences containing nonproteinogenic characters, as well as outliers of the alignment were manually removed from the selection. Sequence headers were annotated according to taxonomy using SeqScrub^64^ and sequence redundancy of >99% was removed. Remaining sequences were aligned using the G-INS-i method of MAFFT and a tree was calculated by FastTree^65^ using the Whelan and Goldman substitution model and standard options for increased accuracy. All sequences forming the clade of the taxonomic group of Metazoa in the tree were extracted and their conserved residues were visualized using WebLogo 3.^63^

### Establishment of NNK-randomized libraries

In a first PCR random mutations in 5UMR were inserted for libraries 5UMR_c (primer 5UMR_c_NNK1), 5UMR_cp (primer 5UMR_cp_NNK1), 5UMR_cy (primer 5UMR_cy_NNK1) with reverse primer 5UMR_PCR1_rev (all primer sequences shown in Suppl. Table 1) with a Q5 HiFi DNA Polymerase (New England Biolabs), followed by preparative Agarose gel purification. 5UMR_wt was amplified using primer 5UMR_PCR2_fwd and primer 5UMR_PCR2_rev. Products from the first PCR were amplified in a subsequent amplification (200 µL reaction volume) using primers 5UMR_cb_NNK2 on PCR product 5UMR_c and primer 5UMR_PCR2_rev for the construction of library 5UMR_cb or primers 5UMR_PCR2_fwd and 5UMR_PCR2_rev for the remaining libraries.

Likewise, primers 3SHU_1_fwd, 3SHU_2_fwd, 3SHU_3_fwd, 3SHU_4_fwd and 3SHU_1_rev (for libraries 3SHU_1, 3SHU_2 and 3SHU_3) or 3SHU_4_rev (for library 3SHU_4) were used for the amplification with Q5® HiFi DNA Polymerase. 3SHU_wt was amplified using primers 3SHU_PCR2_fwd and 3SHU_PCR2_rev, followed by preparative Agarose gel purification and further amplification (200 µl reaction volume) using primers 3SHU_PCR2_fwd and 3SHU_PCR2_rev.

The final library constructs encoded for Aga2p-HA-tag-(G_4_S)_3_ linker-NNK randomized 3SHU or 5UMR gene–c-myc tag. All primers were ordered at Sigma-Aldrich. Amplified PCR2 products were used for ethanol purification and electroporation of *S. cerevisiae* strain EBY100 (ATCC) as described before.^66^ Diversities of the NNK randomized yeast libraries averaged around ∼10^7^ individual clones after electroporation.

Amplified genes for 3SHU_wt and 5UMR_wt were assembled into a BamHI/NheI digested pCTCON2V vector using HiFi DNA Assembly (New England Biolabs). The sequence-verified pCTCON2V plasmid was used for chemical transformation of EBY100 using Frozen-EZ Yeast Transformation II kit (Zymo Research).

### Establishment of final scaffold yeast libraries

Trinucleotide-synthesized primers were ordered from Ella Biotech. 3SHU or 5UMR WT genes were randomized in an initial PCR using primers 3SHU_lib3_PCR1_fwd and 3SHU_lib3_PCR1_rev for the 3SHU gene or 5UMR_cp_PCR1_fwd and 5UMR_PCR1_rev for the 5UMR gene using Q5 HiFi DNA Polymerase (primer sequences shown in Suppl. Table 1; X01 refers to codons encoding for a defined AA frequency as specified in column 2 or 3 in Fig. 2C, while Z01 encodes for the same AA frequency but as reverse codons; X02 refers to codons encoding for an AA frequency for positions Q28 and Q44 in 5UMR, which differs from the remaining positions, Fig. 2C). 10 ng of gel-purified gene fragments of the correct amplicon size were amplified in large volume (200 µL per electroporation) with either primers 3SHU_PCR2_fwd and 3SHU_PCR2_rev or primers 5UMR_PCR2_fwd and 5UMR_PCR2_rev using Q5® HiFi DNA Polymerase. 20 electroporations of EBY100 cells with ethanol-purified DNA were performed as described previously,^66^ reaching a total diversity of 3.6×10^8^ for the 3SHU_final library and 5.3×10^8^ for the 5UMR_final library. Libraries were frozen in SD-CAA+15% glycerol and stored at −80 °C.

### Flow cytometric analysis of yeast libraries

Yeast cultures were cultured in SD-CAA at 30 °C and surface expression was induced in SG-CAA at 20 °C as described previously.^45,67^ Cells were harvested, centrifuged, washed with ice-cold PBSA and resuspended in PBSA to the desired cell concentration. Typically, 1 or 2×10^6^ cells were used for staining in 96-well V-bottom plates with 5 µg/mL anti-c-myc-AF488 antibody (Thermo Fisher Scientific, clone 9E10) and 1 µg/mL anti-HA-AF647 antibody (BioLegend, clone 16B12). Samples were incubated for 30 minutes (4 °C, shaking, in the dark), subsequently washed twice with 200 µL ice-cold PBSA and analyzed with a Cytoflex S instrument (Beckman Coulter).

### Analysis of AA frequency of yeast libraries

DNA of yeast libraries was isolated using an adapted protocol of the Zymoprep Yeast Plasmid Miniprep Kit II (Zymo Research), using 8 µL of Zymolase and an incubation of 3 hours at 37 °C for increased cell wall degradation. DNA isolates were used for the electroporation of 10-beta *E. coli* (New England Biolabs) according to the manufacturer’s protocol using 1 µL of Zymoprep isolate (for 25 µL *E. coli*). After overnight incubation on selective LB agar plates, individual *E. coli* colonies were used for Sanger sequencing in a 96-well plate format (Microsynth). The resulting nucleotide sequences were translated (EMBOSS Transeq, https://www.ebi.ac.uk/Tools/st/emboss_transeq/)^68,69^ and the AA sequence was used for Multiple Sequence Alignment (MSA) (Clustal Omega, https://www.ebi.ac.uk/Tools/msa/clustalo/).^68,70,71^

### Yeast surface display selections

Yeast surface display selections against a variety of targets (CD22, CD276 and an adapter peptide) were carried out. In general, all selections started with a pooled 5UMR/3SHU superlibrary (5UMR_final + 3SHU_final). 10^10^ cells (covering 10x the diversity of the pooled library) were used for the initial rounds of selections. Selection campaigns started with magnetic bead selections using Dynabeads Biotin Binder (Thermo Fisher Scientific) as described previously.^45,66^

Yeast display selections for miniCARbids against CD22 were based on a soluble, biotinylated CD22 protein (AcroBiosystems, SI2-H82E3). Initially, 2 rounds of bead selections were performed with the naïve 5UMR/3SHU superlibrary, followed by a round of error prone PCR (epPCR, GeneMorph II Random Mutagenesis Kit, Agilent) and several rounds of flow cytometric sorting, including a round of negative sorting using a CD276 protein, which carried the same tags (Biotinylated Human B7-H3 (4Ig), His, Avitag; B7B-H82E8).

For yeast display selections against CD276, we used two recombinant proteins by AcroBiosystems (Biotinylated Human B7-H3 (4Ig), B7B-H82E8 and Human B7-H3, B73-H52E2), blocking strategies with anti-CD276 antibody 8H9 (Thermo Fisher Scientific) and cell based pannings with Caco-2 cells. Initially, all sorting strategies started with the naïve 5UMR/3SHU superlibrary and two rounds of bead selections with CD276 4Ig, followed by a round of epPCR and one flow cytometric sort with CD276 4Ig. Subsequently, the yeast library was split up to five different sorting campaigns including selections against CD276 4Ig only (with and without blocking sorts with 8H9), selections incorporating both 2Ig and 4Ig isoforms (with and without blocking sorts with 8H9) and selections incorporating cell pannings^72^ on Caco-2 cells as well as flow cytometric selections with CD276 4Ig.

Yeast display selections for miniCARbids against the adapter peptide (GGGGSYVVERWRHRP) incorporated two rounds of bead selections, in total two rounds of epPCR and eight rounds of flow cytometric sorting. The peptide with different labels (N-terminal FITC or biotin) was used for the selections and synthesized at peptides&elephants.

### Flow cytometric sorting of yeast libraries

For cell sorting either a FACS Aria Fusion cell sorter (BD Biosciences) or SH800S cell sorter (Sony Biotechnology) were used. Either 3×10^7^ or 5×10^6^ yeast cells were harvested and washed twice with 1 mL ice-cold PBSA. All subsequent washing steps were conducted likewise. Yeast cells were stained with the antigen for one hour (4 °C, shaking). The incubation time and staining volume to reach an equilibrium and avoid ligand depletion was calculated as described elsewhere.^67^ Subsequently cells were washed twice and stained with a secondary staining reagent, when required (30 min, 4 °C, shaking, in the dark) with either 5 µg/mL anti-penta-His (conjugated to either AF647 or AF488, Qiagen) or 5 µg/mL streptavidin conjugated to either AF647 or AF488 (Thermo Fisher Scientific), as wells as anti-HA-AF647, anti-HA-AF488 (2 µg/mL, BioLegend, clone 16B12), anti-c-myc-AF488 or anti-c-myc-AF647 (5 µg/mL, Invitrogen, clone 9E10). Prior to sorting, the yeast cells were resuspended in ice-cold PBSA and after sorting incubated in SD-CAA with 100 U/mL penicillin and 100 µg/mL streptomycin at 30 °C while shaking.

### Titration of antigen on yeast displayed miniCARbids

For the titration of peptide-miniCARbids displayed on yeast cells two different antigens were used: biotinylated peptide (Biotin-GGGGSYVVERWRHRP-OH) or a SUMO fusion protein with the adapter peptide sequence at its C-terminus (Suppl. Table 1) with an N-terminal 6xHis-tag for detection. 1×10^6^ PBSA-washed yeast cells (50,000 displaying, 950,000 non-induced cells) were stained in 200 µL overnight (4 °C, shaking, 96-well V-bottom plates). The number of displaying cells, incubation time and staining volume to reach an equilibrium and avoid ligand depletion was calculated as described elsewhere.^67^ Subsequently, samples were washed thrice with 200 µL ice-cold PBSA and stained with 2 µg/mL anti-HA-AF488 and either 5 µg/mL anti-penta-His-AF647 or 20 µg/mL streptavidin-AF647 for 30 minutes (4 °C, shaking, in the dark). After two more washing steps, cells were pelleted and resuspended just before measurement with a Cytoflex S instrument.

### Titration of soluble miniCARbids on cell lines

Binding experiments with CD22-miniCARbids: Cells were harvested and washed with ice-cold PBSA (200 g, 5 min, 4 °C). For affinity determination, 100,000 NALM6 cells were incubated with a range of CD22-miniCARbids concentrations for 4 hours (4 °C, shaking, 96-well V-bottom plates), followed by two washing steps (200 µL ice-cold PBSA) and staining with 5 µg/mL anti-penta-His-AF488 for 20 min (4 °C, shaking, dark) in 25 µL staining volume. After two more washing steps the cells were pelleted and resuspended just before measurement with a Cytoflex S instrument. Data were fitted with a 1:1 binding model to calculate the *K*_D_ values as described previously.^67^ Analysis for unspecific binding was performed likewise, but staining was only performed with 250 nM CD22-miniCARbids for 1 hour (4 °C, shaking).

Binding experiments with CD276-miniCARbids: Titrations with CD276-miniCARbids were performed likewise as described for the analysis of CD22-miniCARbids, but 100,000 cells were stained for 1 hour.

### Quantification of CD22 and CD276 expression on cell lines

Jurkat, NALM6 and Raji cells were harvested and washed twice with ice-cold PBSA (200 g, 5 min, 4 °C). 100,000 cells were stained in 100 µL PBSA with prior blocking (5 min, 4 °C, shaking) of the Fc-receptors using 1 µL TruStain FcX (BioLegend). Subsequently cells were stained with 1 µg/mL anti-CD22-PE antibody (BioLegend, clone HIB22) for 20 min (4 °C, shaking, dark). After two washing steps with 200 µL ice-cold PBSA, cells were resuspended just before measurement with a Cytoflex S instrument. For quantification purposes, Quantibrite PE beads (BD Biosciences) were reconstituted in 500 µL PBSA and analyzed in parallel. Quantification analysis assumed a binding ratio mAb:CD22 of 1:1 and obtained antigen densities were corrected for the fluorophore labelling ratio (PE:mAb) of the antibody.

A-431, A-549, SK-BR-3, Caco-2 and Jurkat cells were analyzed likewise with 3.5 µg/mL of anti-CD276-PE antibody (BioLegend, clone MIH42).

### Blocking experiments of CD276-miniCARbids on Caco-2 cells

100,000 Caco-2 cells were washed with ice-cold PBSA and subsequently blocked with 666 nM of 8H9 or MGA271 (Abeomics) for 1 hour (4°C, shaking, 96-well V-bottom plates) in 18.75 µL staining volume. Subsequently, CD276-miniCARbid was added to reach a final concentration of 500 nM miniCARbid and 500 nM antibody in 25 µL. Samples were incubated for 1 hour (4 °C, shaking, 96-well V-bottom plates) and subsequently washed twice with 200 µL ice-cold PBSA (200 g, 5 minutes, 4 °C). After a secondary staining step with 5 µg/mL anti-penta-His-AF488 for 30 min (4 °C, shaking, dark), two more washing steps were conducted prior to analysis with a Cytoflex S instrument.

### Expression and activation of novel CD22- and CD276-CARs in a Jurkat Nur77 reporter cell line

*In vitro* transcription of CAR transgenes was performed with HiScribe T7 ARCA mRNA Kit, followed by RNA purification with Monarch RNA Cleanup Kit (both from New England Biolabs). Electroporation was performed as described above, using 2×10^6^ Jurkat Nur77 reporter cells. After 2 hours of incubation, cells were counted using a Vi-CELL XR (Beckman Coulter). 50,000 Jurkat Nur77 reporter cells were co-cultured with 100,000 target cells (A-549 for CD276-CARs, NALM6 for CD22-CARs) or no target cells in 200 µL in 96-well U-bottom plates in full-growth medium overnight (15-18 hours). Samples were transferred to 96-well V-bottom plates, washed with 200 µL ice-cold PBSA (300 g, 5 min, 4 °C), resuspended in 50 µL PBSA containing 10% of human serum (PAN Biotech) and incubated for 10 minutes at 4 °C. Subsequently, samples were stained with 2.5 µg/mL anti-MAP-AF647 (BioLegend, clone pmab-1) for 30 min (4 °C, shaking, dark) and finally washed two times prior to analysis with a Cytoflex S instrument.

### Construction of transgenes

The sequence for scFv 376.96 was used in a V_L_-3xG_4_S-V_H_ orientation (patent US 10,233,226 B2); scFv HA22: V_H_-3xG_4_S-V_L_ orientation;^73^ scFv m971 originates from patent US 10,072,078 B2 and was used in a V_H_-linker-V_L_ orientation with both a 1xG_4_S and a 4xG_4_S linker;^27^ scFv MGA271 V_H_-3xG_4_S-V_L_ orientation (patent US 2018/0346544 A1). The 28ζ backbone used was published in Dobersberger et al.,^31^ while the BBζ (Q65K) backbone was published in Salzer et al.^9^

### *In vitro* analysis of peptide-miniCARbids

#### Lentiviral transduction

Peptide-miniCARbids were cloned into a second-generation CAR backbone containing a GM-CSF leader sequence, linker, miniCARbid sequences, 2xG_4_S linker, IgG4 hinge, CD8 transmembrane domain, 4-1BB and CD3ζ. As a benchmark an scFv based binding domain specific for the FGFR2 epitope tag was used in the same backbone, though lacking the miniCARbid-flanking linkers and with an IL22Ra leader sequence. In all constructs, truncated low-affinity nerve growth factor receptor (LNGFR) served as a transduction marker (aa 1-274, Ref sequence: NP_002498.1), which was separated from the CAR via a P2A site. Primary human T cells were isolated from PBMCs of five donors using the Pan T cell isolation Kit, human (Miltenyi Biotec). Subsequently T cells were activated using TransAct (Miltenyi Biotec) according to the manufacturer’s instructions. On the following day, the activated T cells were lentivirally transduced with a MOI of 15 – 20 and washed one day later. The cells were cultivated in TexMACS (Miltenyi Biotec) containing 155 U/mL IL-7 (Miltenyi Biotec) and 290 U/mL IL-15 (Miltenyi Biotec). On day 6, LNGFR positive cells were enriched using the MACSelect™ LNGFR MicroBeads (Miltenyi Biotec) according to the manufacturer’s instructions to 60 – 90% LNGFR+ cells.

#### Cytotoxicity assay and analysis of activation markers

On day 8 – 11 post T cell isolation, 1×10^5^ AdCAR T cells were used for setting up a co-culture with 5×10^4^ CD33+ OCl-AML2 target cells and 50 or 500 ng/mL of CD33-peptide adapter in target cell medium. Target cell lysis was analyzed using flow cytometry on day four post co-culture initiation with a MACSQuant® 10 Analyzer or MACSQuant® X (Miltenyi Biotec). For this, 100 µL of cells were transferred into a new 96 well plate and 2×10^3^ CountBright absolute counting beads (Thermo Fisher Scientific) were added. After centrifugation and removal of supernatant, T cells were stained in 50 µL CliniMACS® PBS/ EDTA buffer supplemented with 0.5% BSA (Miltenyi Biotec) and following antibodies (all Miltenyi Biotec): 7AAD staining solution, CD3-Vioblue (REA613), CD271 (LNGFR)-Viogreen (REA844), CD25-PE-Vio770 (REA570), CD137-APC (REA765) and CD69-APC-Vio770 (REA824) for 10 min at 4°C. All antibodies were used according to manufacturer’s protocol. After washing, cells were analyzed by flow cytometry and cell counts were calculated based on analyzed CountBright absolute counting bead numbers.

#### Cytokine release by CAR T cells

Release of cytokines IFN-Υ, TNF-α, GM-CSF and IL-2 were analyzed one day after the initiation of co-culture with 50 ng/mL CD33-peptide adapter using the MACSPlex Cytokine 12 Kit, human (all Miltenyi Biotec) according to the manufacturer’s protocol.

### *In vitro* analysis of CD22 and CD276-miniCARbids

#### Lentiviral transduction

Lenti-X 293T cells (Takara) were split one day prior to transfection in DMEM (Thermo Fisher Scientific) + 10% FCS (Capricorn Scientific) to reach confluency of 70-80% at the time of transfection. Lenti-X 293T cells were co-transfected with a pCDH expression vector (System Biosciences) carrying puromycin resistance genes and second-generation viral packaging plasmids pMD2.G and psPAX2 (Addgene plasmids #12259 and #12260, respectively; gifts from Didier Trono) using the PureFection Transfection Reagent (System Biosciences). The transfection mixture was vortexed for 10 seconds and incubated at RT for 15 min, before the dropwise addition to the cells. Supernatants were collected on day two and three after transfection and concentrated 50x using Lenti-X Concentrator (Takara). Lentiviral particles were resuspended gently in AIM V (Thermo Fisher Scientific) supplemented with 2% Octaplas (Octapharma) supplemented with 2.5% HEPES (PAN-Biotech), 1% L-Glutamine (Gibco) and 200 U/ml IL-2 (Peproptech) and stored at −80 °C. One day prior to transduction primary human T cells were activated with T-Activator CD3/CD28 dynabeads (Thermo Fisher Scientfic) according to the manufacturer’s instructions in AIM V™ medium supplemented with 2% Octaplas, 2.5% HEPES, 1% L-Glutamine and 200 U/ml IL-2. Activated T cells (10^6^ cells/mL) were transduced in retronectin (Takara) coated cell culture dishes with thawed lentiviral suspension at a final dilution of 1:2. To ensure growth of transduced cells only, puromycin was added to a final concentration of 1 µg/mL for 48 hours two days post transduction. Transduced T cells were expanded at densities below 1.5×10^6^ cells/mL to ensure high cell fitness and stained for the expression of CD3 (anti-CD3-Viogreen, Miltenyi, clone REA613, final dilution 1:50), CD4 (anti-CD4-PerCP, BioLegend, clone OKT4, 1:30), CD8 (anti-CD8-FITC, Miltenyi, clone REA734, 1:50) and the CAR (anti-MAP-PE, Novus Bio, clone pmab-1, 1:250), incubated for 25 min, washed twice and analyzed with a BD FACSymphony™ A5 Cell Analyzer instrument (BD Biosciences).

#### Cytokine release by CAR T cells

Cytokine secretion was quantified in the supernatants from cytotoxicity experiments (described above). Supernatants were centrifuged (500 g, 5 min, 4 °C) and subsequently stored at −20 °C. IFN-γ and IL-2 were quantified using ELISA MAX Deluxe Set for human IFN-γ and IL-2 (Biozym).

#### Cytotoxicity assay

A luciferase-assay was used to determine the cytotoxicity of primary human T cells. For anti-CD22 CAR T cells GFP & luciferase expressing target cell lines NALM6 and Raji were co-cultured at E:T ratios 2:1 and 5:1 for either 4 h (NALM6) or 24 h (Raji) in 100 µL RPMI without phenol red (Thermo Fisher Scientific, RPMI 1640 Medium, no phenol red) supplemented with 2% Octaplas and 1% penicillin-streptomycin (Thermo Fisher Scientific) in white 96-well U-bottom plates. For anti-CD276 CAR T cells GFP & luciferase expressing target cells lines A-431, A-549 and Caco-2 were seeded 16 h prior to the assay in 100 µL RPMI without phenol red supplemented with 2% Octaplas and 1% penicillin-streptomycin in white round-bottom 96-well plates. On the following day 50 µL primary T cells were added to reach E:T ratios of 5:1 and 10:1 for either 6 or 24 h. After co-culture the number of viable cells was determined by their luciferase activity. The plates were equilibrated 10 min at RT, before luciferin (Revvity) was added to reach a final concentration of 150 µg/mL. Luminescence was quantified after 20 min at RT using an ENSPIRE Multimode plate reader (Perkin Elmer). The lysis of target cells was calculated as following:

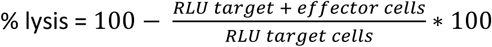

### *In vivo* experiments

NOD.Cg-Prkdcscid Il2rgtm1Wjl/SzJ (NSG, The Jackson Laboratory) mice were bred at the core facility laboratory animal breeding and husbandry of the Medical University of Vienna under specific pathogen-free conditions according to FELASA recommendations. All practices were approved by the Ethics and Animal Welfare Committee of the Medical University of Vienna and granted by the national authority (Austrian Federal Ministry of Education, Science and Research) according to Section 8ff of the Law for Animal Experiments under license GZ 66.009/0243-V/3b/2019 and were performed according to the guidelines of FELASA and ARRIVE. Primary human T cells were lentivirally transduced to express a m971-1xG_4_S-, 22_1410- or 22_1333-CAR and expanded for 13 days. 0.5×10^5^ NALM6 cells in 200 µL RPMI were injected i.v. into the tail veins of NSG mice (female, 10-13 weeks of age). Three days later 5×10^6^ CAR T cells or untransduced T cells (“mock”) in 200 µL RPMI were injected i.v. into the tail veins likewise. The mice were monitored daily and sacrificed nine days after the injection of CAR T cells.

Bone marrow single-cells suspensions were obtained by flushing two femurs with 20 mL PBS and subsequent filtering through a 70 µm cell strainer. Spleen samples were disrupted and passed through 70 µm filters twice, while the liver samples were filtered once following disruption. Liver single-cell suspensions were subjected to a 33.75% Percoll gradient centrifugation to separate lymphocytes from hepatocytes. All single-cell suspensions were subjected to multiple washing steps and lysis of red blood cells through ammonium chloride potassium (ACK) Lysing Buffer. Finally, cells were blocked with 5% Octaplas and a fourth of the volume was stained in 50 µL antibody mix (anti-CD3-BV605, BioLegend, clone OKT3, final dilution 1:50; anti-MAP-PE, Novus Bio, clone pmab-1, 1:100; anti-CD19-APC, Miltenyi Biotec, clone REA675, 1:100; fixable viability dye eF780, Invitrogen, 1:1000), washed twice and measured on a BD FACSymphony™ A3 Cell Analyzer instrument together with 10 µL beads.

## References

1 Cappell, K. M. & Kochenderfer, J. N. Long-term outcomes following CAR T cell therapy: what we know so far. Nat Rev Clin Oncol 20, 359–371 (2023). 10.1038/s41571-023-00754-1

2 June, C. H. & Sadelain, M. Chimeric Antigen Receptor Therapy. N Engl J Med 379, 64–73 (2018). 10.1056/NEJMra1706169

3 Labanieh, L. & Mackall, C. L. CAR immune cells: design principles, resistance and the next generation. Nature 614, 635–648 (2023). 10.1038/s41586-023-05707-3

4 Majzner, R. G. & Mackall, C. L. Clinical lessons learned from the first leg of the CAR T cell journey. Nat Med 25, 1341–1355 (2019). 10.1038/s41591-019-0564-6

5 Ajina, A. & Maher, J. Strategies to Address Chimeric Antigen Receptor Tonic Signaling. Mol Cancer Ther 17, 1795–1815 (2018). 10.1158/1535-7163.MCT-17-1097

6 Long, A. H. et al. 4-1BB costimulation ameliorates T cell exhaustion induced by tonic signaling of chimeric antigen receptors. Nat Med 21, 581–590 (2015). 10.1038/nm.3838

7 Arndt, K. M., Muller, K. M. & Pluckthun, A. Factors influencing the dimer to monomer transition of an antibody single-chain Fv fragment. Biochemistry 37, 12918–12926 (1998). 10.1021/bi9810407

8 Todorovska, A., Roovers, R. C., Dolezal, O., Kortt, A. A., Hoogenboom, H. R. & Hudson, P. J. Design and application of diabodies, triabodies and tetrabodies for cancer targeting. J Immunol Methods 248, 47–66 (2001). 10.1016/s0022-1759(00)00342-2

9 Salzer, B. et al. Engineering AvidCARs for combinatorial antigen recognition and reversible control of CAR function. Nat Commun 11, 4166 (2020). 10.1038/s41467-020-17970-3

10 Qin, H. et al. Preclinical Development of Bivalent Chimeric Antigen Receptors Targeting Both CD19 and CD22. Mol Ther Oncolytics 11, 127–137 (2018). 10.1016/j.omto.2018.10.006

11 Counsell, J. R. et al. Lentiviral vectors can be used for full-length dystrophin gene therapy. Sci Rep 7, 44775 (2017). 10.1038/srep44775

12 Sweeney, N. P. & Vink, C. A. The impact of lentiviral vector genome size and producer cell genomic to gag-pol mRNA ratios on packaging efficiency and titre. Mol Ther Methods Clin Dev 21, 574–584 (2021). 10.1016/j.omtm.2021.04.007

13 Zhu, I. et al. Modular design of synthetic receptors for programmed gene regulation in cell therapies. Cell 185, 1431–1443 e1416 (2022). 10.1016/j.cell.2022.03.023

14 Turtle, C. J. et al. CD19 CAR-T cells of defined CD4+:CD8+ composition in adult B cell ALL patients. J Clin Invest 126, 2123–2138 (2016). 10.1172/JCI85309

15 Wagner, D. L. et al. Immunogenicity of CAR T cells in cancer therapy. Nat Rev Clin Oncol 18, 379–393 (2021). 10.1038/s41571-021-00476-2

16 De Munter, S. et al. Nanobody Based Dual Specific CARs. Int J Mol Sci 19 (2018). 10.3390/ijms19020403

17 Hammill, J. A. et al. Designed ankyrin repeat proteins are effective targeting elements for chimeric antigen receptors. J Immunother Cancer 3, 55 (2015). 10.1186/s40425-015-0099-4

18 Han, X., Cinay, G. E., Zhao, Y., Guo, Y., Zhang, X. & Wang, P. Adnectin-Based Design of Chimeric Antigen Receptor for T Cell Engineering. Mol Ther 25, 2466–2476 (2017). 10.1016/j.ymthe.2017.07.009

19 Siegler, E., Li, S., Kim, Y. J. & Wang, P. Designed Ankyrin Repeat Proteins as Her2 Targeting Domains in Chimeric Antigen Receptor-Engineered T Cells. Hum Gene Ther 28, 726–736 (2017). 10.1089/hum.2017.021

20 Zajc, C. U. et al. A conformation-specific ON-switch for controlling CAR T cells with an orally available drug. Proc Natl Acad Sci U S A 117, 14926–14935 (2020). 10.1073/pnas.1911154117

21 Zajc, C. U., Salzer, B., Taft, J. M., Reddy, S. T., Lehner, M. & Traxlmayr, M. W. Driving CARs with alternative navigation tools - the potential of engineered binding scaffolds. FEBS J 288, 2103–2118 (2021). 10.1111/febs.15523

22 Julian, M. C. et al. Co-evolution of affinity and stability of grafted amyloid-motif domain antibodies. Protein Eng Des Sel 28, 339–350 (2015). 10.1093/protein/gzv050

23 Julian, M. C., Li, L., Garde, S., Wilen, R. & Tessier, P. M. Efficient affinity maturation of antibody variable domains requires co-selection of compensatory mutations to maintain thermodynamic stability. Sci Rep 7, 45259 (2017). 10.1038/srep45259

24 Porebski, B. T. et al. Circumventing the stability-function trade-off in an engineered FN3 domain. Protein Eng Des Sel 29, 541–550 (2016). 10.1093/protein/gzw046

25 Teufl, M., Zajc, C. U. & Traxlmayr, M. W. Engineering Strategies to Overcome the Stability-Function Trade-Off in Proteins. ACS Synth Biol 11, 1030–1039 (2022). 10.1021/acssynbio.1c00512

26 Haso, W. et al. Anti-CD22-chimeric antigen receptors targeting B-cell precursor acute lymphoblastic leukemia. Blood 121, 1165–1174 (2013). 10.1182/blood-2012-06-438002

27 Singh, N. et al. Antigen-independent activation enhances the efficacy of 4-1BB-costimulated CD22 CAR T cells. Nat Med 27, 842–850 (2021). 10.1038/s41591-021-01326-5

28 Majzner, R. G. et al. CAR T Cells Targeting B7-H3, a Pan-Cancer Antigen, Demonstrate Potent Preclinical Activity Against Pediatric Solid Tumors and Brain Tumors. Clin Cancer Res 25, 2560–2574 (2019). 10.1158/1078-0432.CCR-18-0432

29 Arndt, C., Fasslrinner, F., Loureiro, L. R., Koristka, S., Feldmann, A. & Bachmann, M. Adaptor CAR Platforms-Next Generation of T Cell-Based Cancer Immunotherapy. Cancers (Basel) 12 (2020). 10.3390/cancers12051302

30 Seitz, C. M. et al. Novel adapter CAR-T cell technology for precisely controllable multiplex cancer targeting. Oncoimmunology 10, 2003532 (2021). 10.1080/2162402X.2021.2003532

31 Dobersberger, M. et al. An engineering strategy to target activated EGFR with CAR T cells. Cell Rep Methods 4, 100728 (2024). 10.1016/j.crmeth.2024.100728

32 Gera, N., Hussain, M., Wright, R. C. & Rao, B. M. Highly stable binding proteins derived from the hyperthermophilic Sso7d scaffold. J Mol Biol 409, 601–616 (2011). 10.1016/j.jmb.2011.04.020

33 Mouratou, B. et al. Remodeling a DNA-binding protein as a specific in vivo inhibitor of bacterial secretin PulD. Proc Natl Acad Sci U S A 104, 17983–17988 (2007). 10.1073/pnas.0702963104

34 Paumann-Page, M. et al. Peroxidasin protein expression and enzymatic activity in metastatic melanoma cell lines are associated with invasive potential. Redox Biol 46, 102090 (2021). 10.1016/j.redox.2021.102090

35 Traxlmayr, M. W. et al. Strong Enrichment of Aromatic Residues in Binding Sites from a Charge-neutralized Hyperthermostable Sso7d Scaffold Library. J Biol Chem 291, 22496–22508 (2016). 10.1074/jbc.M116.741314

36 Casadevall, N. et al. Pure red-cell aplasia and antierythropoietin antibodies in patients treated with recombinant erythropoietin. N Engl J Med 346, 469–475 (2002). 10.1056/NEJMoa011931

37 Saxton, R. A., Glassman, C. R. & Garcia, K. C. Emerging principles of cytokine pharmacology and therapeutics. Nat Rev Drug Discov 22, 21–37 (2023). 10.1038/s41573-022-00557-6

38 Hackel, B. J., Ackerman, M. E., Howland, S. W. & Wittrup, K. D. Stability and CDR composition biases enrich binder functionality landscapes. J Mol Biol 401, 84–96 (2010). 10.1016/j.jmb.2010.06.004

39 Koide, A., Bailey, C. W., Huang, X. & Koide, S. The fibronectin type III domain as a scaffold for novel binding proteins. J Mol Biol 284, 1141–1151 (1998). 10.1006/jmbi.1998.2238

40 Lynn, R. C. et al. c-Jun overexpression in CAR T cells induces exhaustion resistance. Nature 576, 293–300 (2019). 10.1038/s41586-019-1805-z

41 Shusta, E. V., Holler, P. D., Kieke, M. C., Kranz, D. M. & Wittrup, K. D. Directed evolution of a stable scaffold for T-cell receptor engineering. Nat Biotechnol 18, 754–759 (2000). 10.1038/77325

42 Traxlmayr, M. W. & Obinger, C. Directed evolution of proteins for increased stability and expression using yeast display. Arch Biochem Biophys 526, 174–180 (2012). 10.1016/j.abb.2012.04.022

43 Bogan, A. A. & Thorn, K. S. Anatomy of hot spots in protein interfaces. J Mol Biol 280, 1–9 (1998). 10.1006/jmbi.1998.1843

44 Zemlin, M. et al. Expressed murine and human CDR-H3 intervals of equal length exhibit distinct repertoires that differ in their amino acid composition and predicted range of structures. J Mol Biol 334, 733–749 (2003). 10.1016/j.jmb.2003.10.007

45 Angelini, A. et al. Protein Engineering and Selection Using Yeast Surface Display. Methods Mol Biol 1319, 3–36 (2015). 10.1007/978-1-4939-2748-7_1

46 Boder, E. T. & Wittrup, K. D. Yeast surface display for screening combinatorial polypeptide libraries. Nat Biotechnol 15, 553–557 (1997). 10.1038/nbt0697-553

47 Bottino, C., Vitale, C., Dondero, A. & Castriconi, R. B7-H3 in Pediatric Tumors: Far beyond Neuroblastoma. Cancers (Basel) 15 (2023). 10.3390/cancers15133279

48 Zhou, W. T. & Jin, W. L. B7-H3/CD276: An Emerging Cancer Immunotherapy. Front Immunol 12, 701006 (2021). 10.3389/fimmu.2021.701006

49 Wang, Z., Yang, J., Zhu, Y., Zhu, Y., Zhang, B. & Zhou, Y. Differential expression of 2IgB7-H3 and 4IgB7-H3 in cancer cell lines and glioma tissues. Oncol Lett 10, 2204–2208 (2015). 10.3892/ol.2015.3611

50 Feustel, K., Martin, J. & Falchook, G. S. B7-H3 Inhibitors in Oncology Clinical Trials: A Review. J Immunother Precis Oncol 7, 53–66 (2024). 10.36401/JIPO-23-18

51 Shah, N. N. et al. CD4/CD8 T-Cell Selection Affects Chimeric Antigen Receptor (CAR) T-Cell Potency and Toxicity: Updated Results From a Phase I Anti-CD22 CAR T-Cell Trial. J Clin Oncol 38, 1938–1950 (2020). 10.1200/JCO.19.03279

52 Ashouri, J. F. & Weiss, A. Endogenous Nur77 Is a Specific Indicator of Antigen Receptor Signaling in Human T and B Cells. J Immunol 198, 657–668 (2017). 10.4049/jimmunol.1601301

53 Smith, E. L. et al. GPRC5D is a target for the immunotherapy of multiple myeloma with rationally designed CAR T cells. Sci Transl Med 11 (2019). 10.1126/scitranslmed.aau7746

54 Fry, T. J. et al. CD22-targeted CAR T cells induce remission in B-ALL that is naive or resistant to CD19-targeted CAR immunotherapy. Nat Med 24, 20–28 (2018). 10.1038/nm.4441

55 Kokalaki, E. et al. Dual targeting of CD19 and CD22 against B-ALL using a novel high-sensitivity aCD22 CAR. Mol Ther 31, 2089–2104 (2023). 10.1016/j.ymthe.2023.03.020

56 Du, H. et al. Antitumor Responses in the Absence of Toxicity in Solid Tumors by Targeting B7-H3 via Chimeric Antigen Receptor T Cells. Cancer Cell 35, 221–237 e228 (2019). 10.1016/j.ccell.2019.01.002

57 Theruvath, J. et al. Locoregionally administered B7-H3-targeted CAR T cells for treatment of atypical teratoid/rhabdoid tumors. Nat Med 26, 712–719 (2020). 10.1038/s41591-020-0821-8

58 Traxlmayr, M. W. et al. Directed evolution of Her2/neu-binding IgG1-Fc for improved stability and resistance to aggregation by using yeast surface display. Protein Eng Des Sel 26, 255–265 (2013). 10.1093/protein/gzs102

59 Frank, M. J. et al. CD22-CAR T-Cell Therapy Mediates High Durable Remission Rates in Adults with Large B-Cell Lymphoma Who Have Relapsed after CD19-CAR T-Cell Therapy. Blood 138 (2021). 10.1182/blood-2021-152145

60 Zallot, R., Oberg, N. & Gerlt, J. A. The EFI Web Resource for Genomic Enzymology Tools: Leveraging Protein, Genome, and Metagenome Databases to Discover Novel Enzymes and Metabolic Pathways. Biochemistry 58, 4169–4182 (2019). 10.1021/acs.biochem.9b00735

61 Katoh, K. & Standley, D. M. MAFFT multiple sequence alignment software version 7: improvements in performance and usability. Mol Biol Evol 30, 772–780 (2013). 10.1093/molbev/mst010

62 Capella-Gutierrez, S., Silla-Martinez, J. M. & Gabaldon, T. trimAl: a tool for automated alignment trimming in large-scale phylogenetic analyses. Bioinformatics 25, 1972–1973 (2009). 10.1093/bioinformatics/btp348

63 Crooks, G. E., Hon, G., Chandonia, J. M. & Brenner, S. E. WebLogo: a sequence logo generator. Genome Res 14, 1188–1190 (2004). 10.1101/gr.849004

64 Foley, G., Sutzl, L., D’Cunha, S. A., Gillam, E. M. & Boden, M. SeqScrub: a web tool for automatic cleaning and annotation of FASTA file headers for bioinformatic applications. Biotechniques 67, 50–54 (2019). 10.2144/btn-2018-0188

65 Price, M. N., Dehal, P. S. & Arkin, A. P. FastTree: computing large minimum evolution trees with profiles instead of a distance matrix. Mol Biol Evol 26, 1641–1650 (2009). 10.1093/molbev/msp077

66 Chen, T. F., de Picciotto, S., Hackel, B. J. & Wittrup, K. D. Engineering fibronectin-based binding proteins by yeast surface display. Methods Enzymol 523, 303–326 (2013). 10.1016/B978-0-12-394292-0.00014-X

67 Zajc, C. U., Teufl, M. & Traxlmayr, M. W. Affinity and Stability Analysis of Yeast Displayed Proteins. Methods Mol Biol 2491, 155–173 (2022). 10.1007/978-1-0716-2285-8_9

68 Goujon, M. et al. A new bioinformatics analysis tools framework at EMBL-EBI. Nucleic Acids Research 38, W695–W699 (2010). 10.1093/nar/gkq313

69 Rice, P., Longden, I. & Bleasby, A. EMBOSS: the European Molecular Biology Open Software Suite. Trends Genet 16, 276–277 (2000). 10.1016/s0168-9525(00)02024-2

70 McWilliam, H. et al. Analysis Tool Web Services from the EMBL-EBI. Nucleic Acids Res 41, W597–600 (2013). 10.1093/nar/gkt376

71 Sievers, F. et al. Fast, scalable generation of high-quality protein multiple sequence alignments using Clustal Omega. Mol Syst Biol 7, 539 (2011). 10.1038/msb.2011.75

72 Panton, R. A. & Stern, L. A. Ligand Selection by Combination of Recombinant and Cell Panning Selection Techniques. Methods Mol Biol 2491, 217–233 (2022). 10.1007/978-1-0716-2285-8_12

73 Ho, M., Kreitman, R. J., Onda, M. & Pastan, I. In vitro antibody evolution targeting germline hot spots to increase activity of an anti-CD22 immunotoxin. J Biol Chem 280, 607–617 (2005). 10.1074/jbc.M409783200

