## Supplementary Information for "MiniCARbids: Minimalistic human binding domains specifically tailored to CAR T applications"

Fig. S1

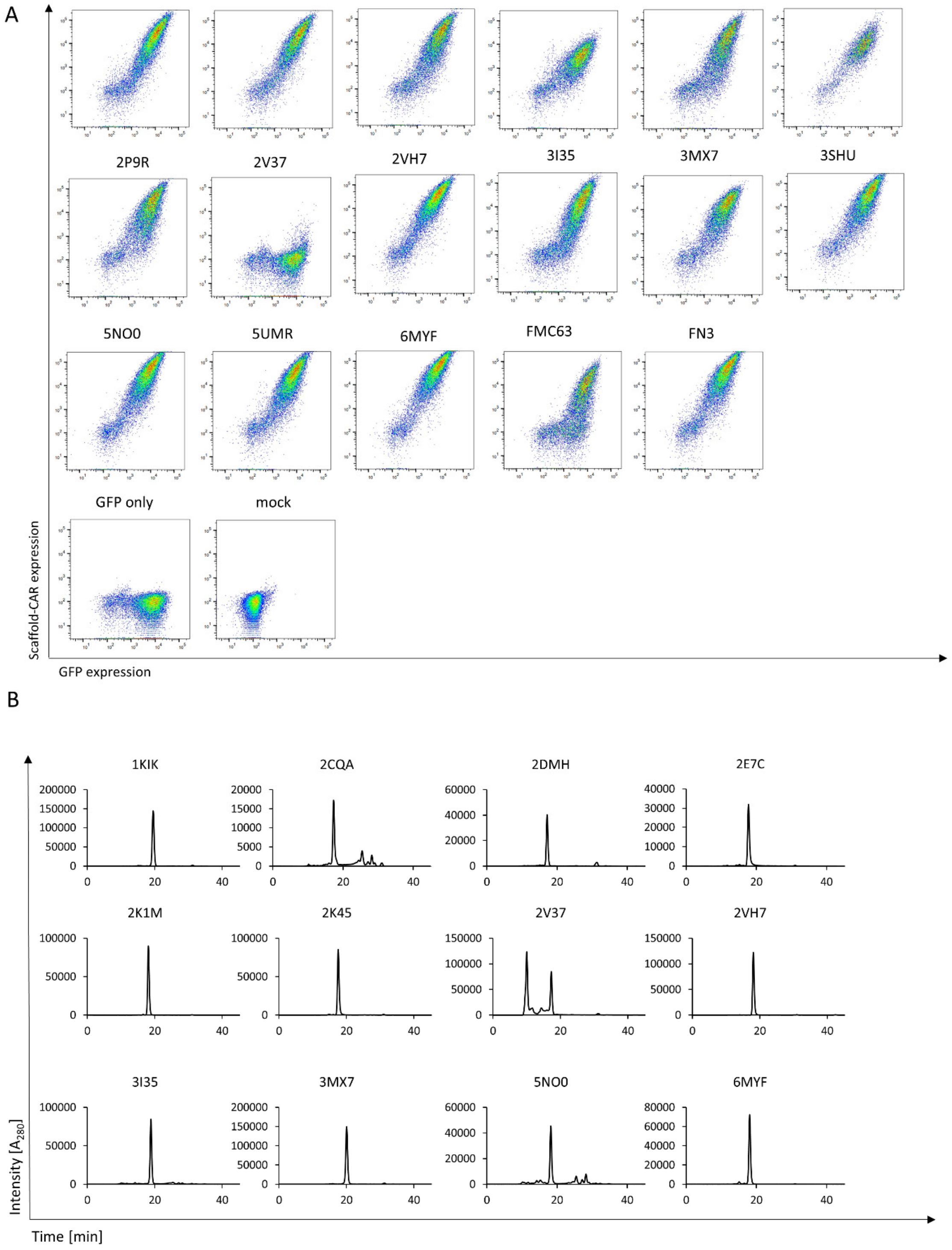


**Suppl. Fig. 1: Assessment of scaffold-CAR expression levels and SEC-HPLC analysis.** (A) Flow cytometric analysis of the expression of 15 scaffold-CARs, FMC63-CAR and FN3-CAR in a BBζ (Q65K) backbone on primary human T cells. Scaffold-CAR mRNA was co-electroporated with GFP mRNA to confirm successful electroporation. The expression level was evaluated via anti-Flag-tag staining by flow cytometry. One representative example is shown (n=4, biological replicates). (B) Aggregation properties were assessed using SEC-HPLC. Chromatograms from one representative experiment (n=3, technical replicates) are shown for the remaining 12 scaffolds (3SHU, 5UMR and 2P9R shown in Fig. 1F), which had been tested for expression on a CAR backbone on primary human T cells. Here 500 µg of scaffold 2V37 was analyzed, yielding comparable results to prior analysis of 50 µg protein.

Fig. S2

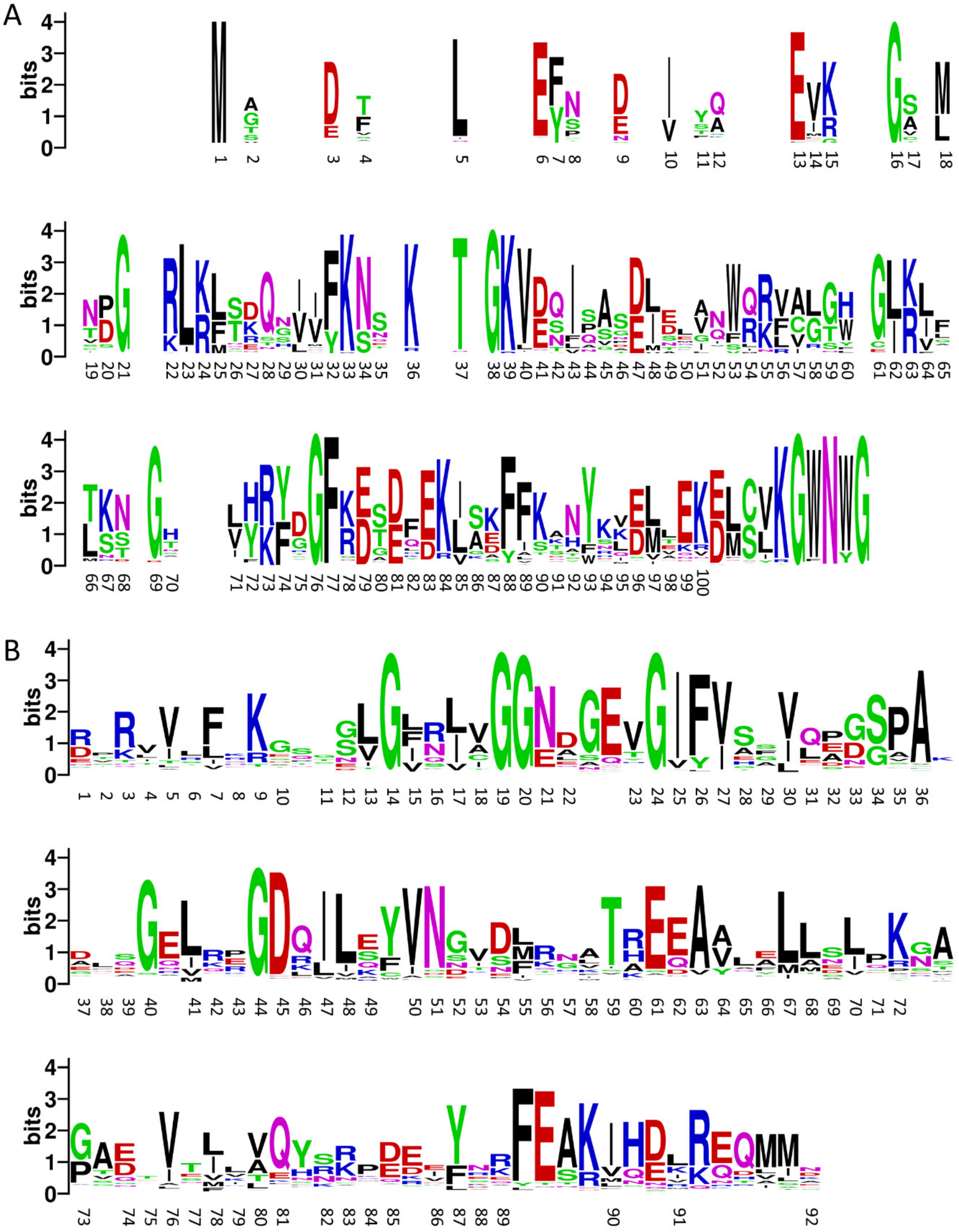

**Suppl. Fig. 2: Phylogenetic analysis of 5UMR and 3SHU to predict conserved residues.** (A) *H. sapiens* sequence of FACT complex subunit SSRP1 and its homolog from *S. cerevisiae* were used for a PSI-BLAST search. Sequences were aligned with MAFFT and sequences forming the clade Metazoa are visualized here. (B) *H. sapiens* sequence of the tight junction protein ZO-1 was used for a UniRef90 search. Sequences were aligned with MAFFT and the refined alignment is visualized here.

Fig. S3

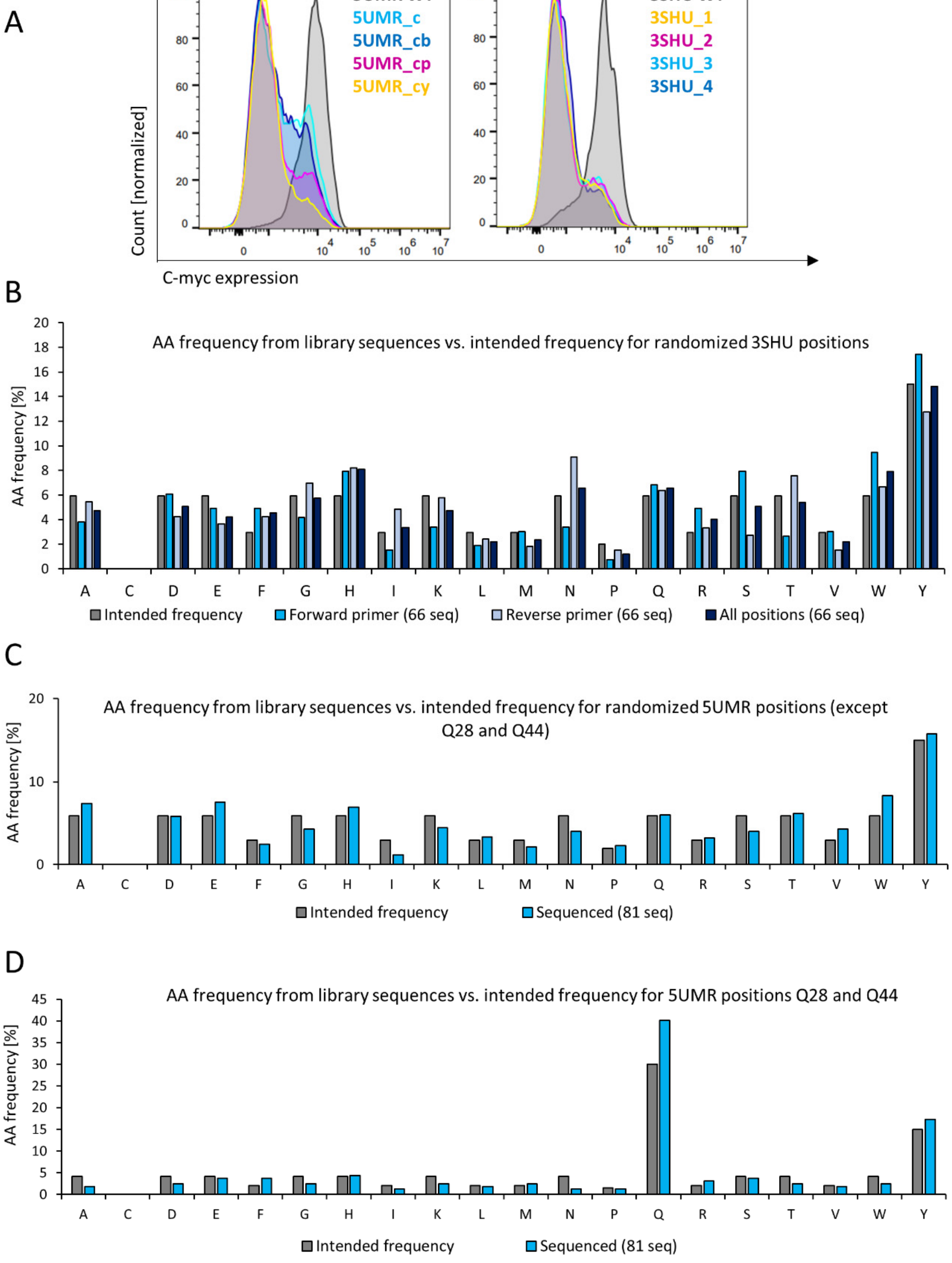


**Suppl. Fig. 3: Analysis of randomly mutated yeast display libraries.** (A) Histograms of c-myc expression – representing full-length expression - of NNK-randomized libraries and their non-mutated parental proteins 3SHU or 5UMR WT. (B) Sequencing results of randomized positions of 66 clones for the assessment of the amino acid frequency in the yeast library 3SHU_final compared to the intended frequency. (C) Sequencing results of all randomized positions in the library 5UMR_final (except Q28 and Q44) of 81 clones for the assessment of the amino acid frequency in the final yeast library compared to the intended frequency. (D) Sequencing results of randomized positions Q28 and Q44 of 81 clones in the library 5UMR_final for the assessment of the amino acid frequency in the final yeast library compared to the intended frequency.

Fig. S4

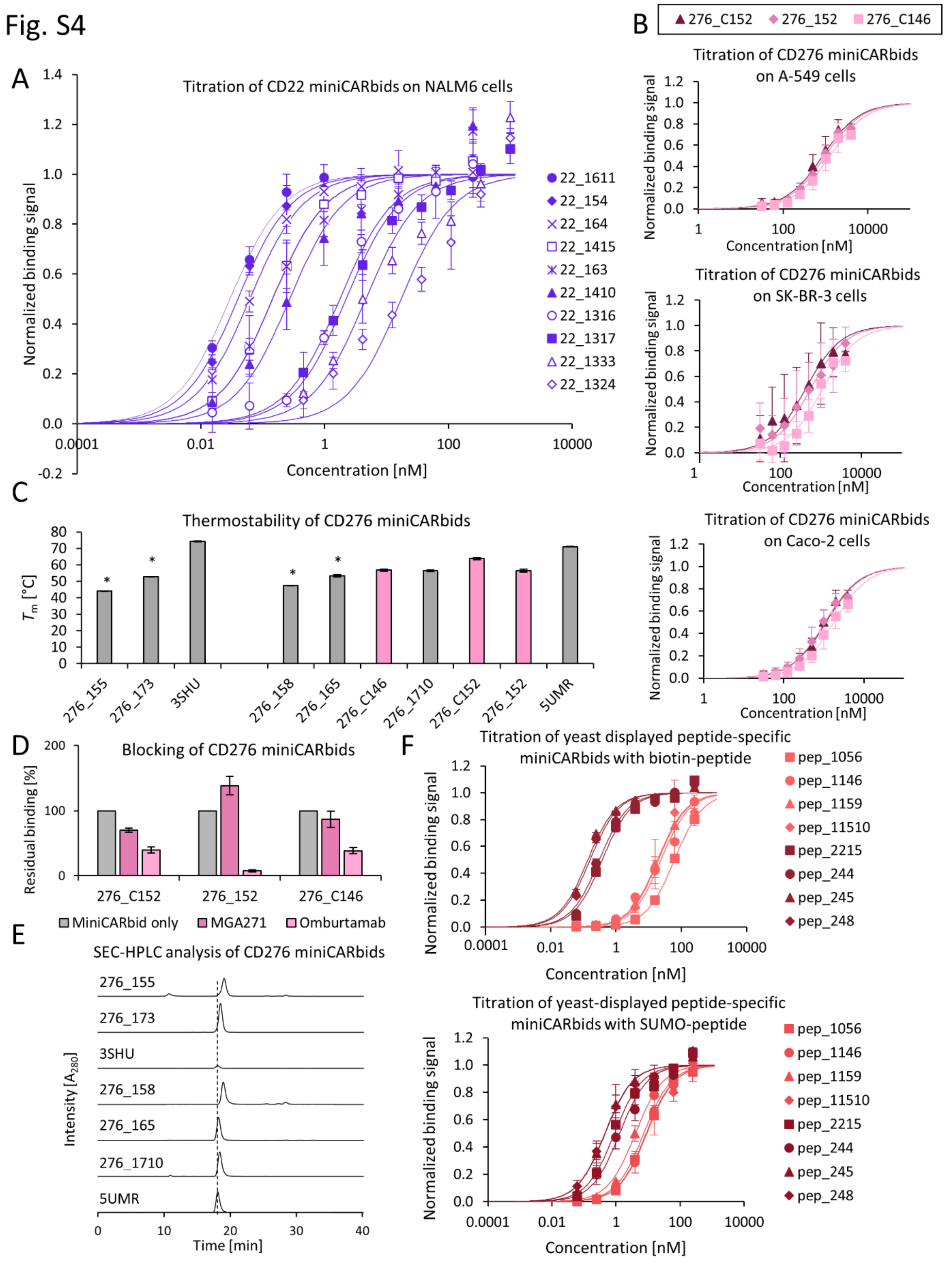


**Suppl. Fig. 4: Detailed biochemical analysis of engineered miniCARbids.** (A) Titrations of ten CD22-miniCARbids on NALM6 are shown (average ± SD, n=3 or 4, biological replicates). The binding intensity was analyzed via anti-His-tag staining by flow cytometry. (B) Titration curves of three CD276-miniCARbids (276_C152, 276_152 and 276_C146) on cell lines A-549 (14,140 ± 940 CD276 molecules/cell, average ± SD, n=3, biological replicates), Caco-2 (45,200 ± 24,290 CD276 molecules/cell, average ± SD, n=3, biological replicates) and SK-BR-3 (2,040 ± 90 CD276 molecules/cell, average ± SD, n=3, biological replicates). The binding intensity was assessed via anti-His-tag staining by flow cytometry. Data were fitted with a 1:1 binding model (solid lines) for the calculation of the respective $K_D$ values (average ± SD, n=3 or 4, biological replicates). (C) $T_m$ values of CD276-miniCARbids as determined by DSC (average ± SD from two or three independent experiments, technical replicates); $T_m$ values from only two independent measurements are marked with an asterisk. (D) Blocking experiments were performed with known anti-CD276 antibodies omburtamab and MGA271. Cells were pre-incubated with 500 nM of omburtamab or MGA271 and subsequently stained with an equimolar concentration of CD276 binder. Detection of CD276-miniCARbids was performed via anti-His-tag staining by flow cytometry (average ± SD, n=3, biological replicates). (E) SEC-HPLC analysis of five additional CD276-miniCARbids and their respective parental proteins 3SHU and 5UMR. One representative example of three independent measurements (technical replicates) is shown. (F) Titrations of yeast-displayed peptide-miniCARbids were performed with both SUMO-peptide and biotin-peptide as antigens. The binding intensity was analyzed via anti-His-tag staining (for SUMO-peptide) or fluorescently labeled streptavidin (for biotin-peptide) by flow cytometry (average ± SD, n=3, biological replicates).

Fig. S5

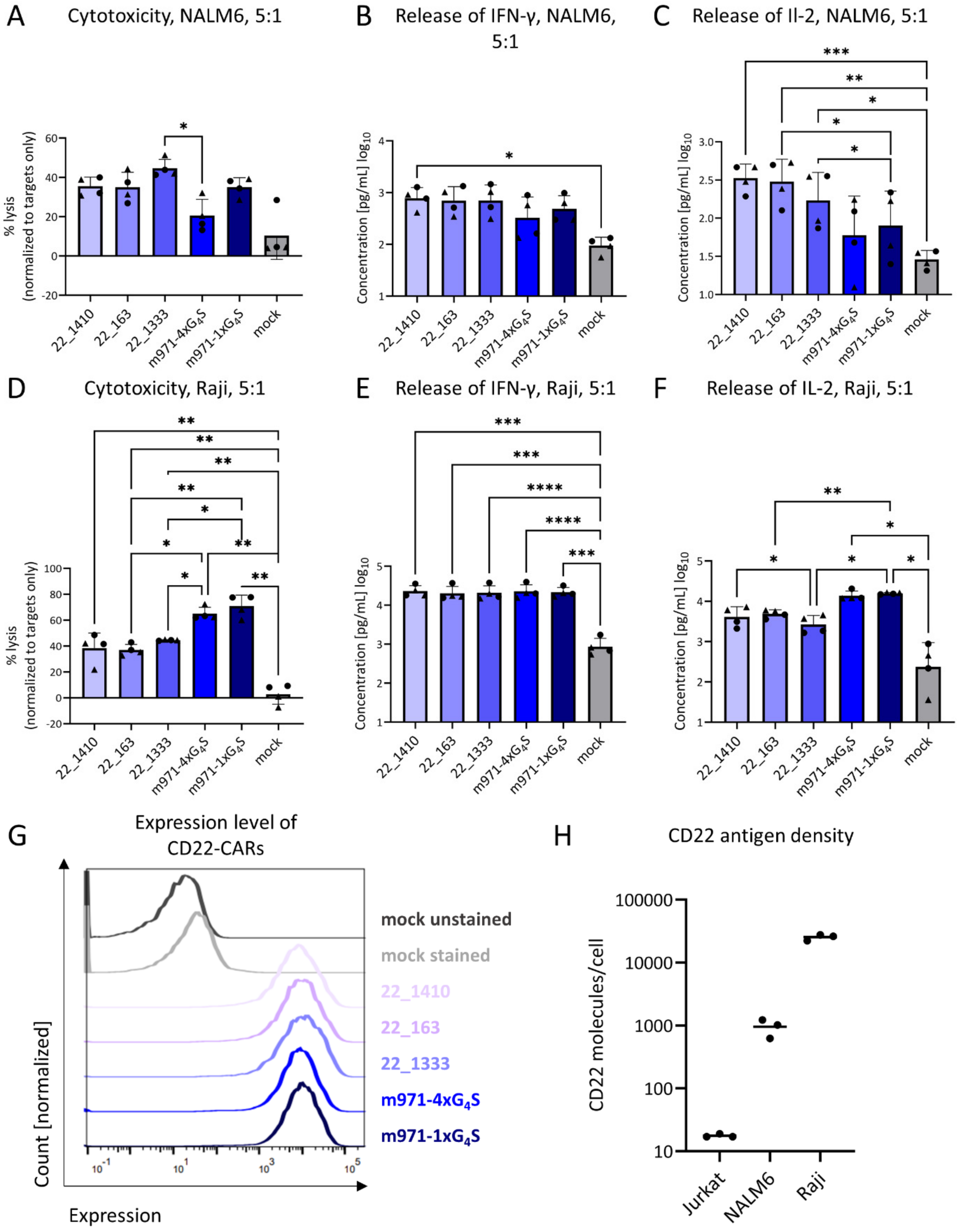


**Suppl. Figure 5: Functional analysis of CD22-specific miniCARbid-CAR Ts *in vitro*.** (A) Cytotoxicity of CD22-specific CAR T cells and mock T cells (no CAR) against NALM6 cells (E:T 5:1, average ± SD, n=4, biological replicates). (B) Release of IFN-γ analyzed via ELISA. The cytokines were analyzed in the supernatants of co-cultures with NALM6 cells (E:T 5:1, average ± SD, n=4, biological replicates). (C) Release of IL-2 analyzed via ELISA. The cytokines were analyzed in the supernatants of co-cultures with

NALM6 cells (E:T 5:1, average ± SD, n=4, biological replicates). (D) Cytotoxicity of CD22- specific CAR T cells and mock T cells (no CAR) against Raji cells (E:T 5:1, average ± SD, n=4, biological replicates). (E) Release of IFN-γ analyzed via ELISA. The cytokines were analyzed in the supernatants of co-cultures with Raji cells (E:T 5:1, average ± SD, n=4, biological replicates). (F) Release of IL-2 analyzed via ELISA. The cytokines were analyzed in the supernatants of co-cultures with Raji cells (E:T 5:1, average ± SD, n=4, biological replicates). (G) Expression levels of CD22-specific CARs on transduced primary T cells as determined by anti-MAP-tag staining and measured by flow cytometry. (H) CD22 antigen density (CD22 molecules/cell) of Jurkat, NALM6 and Raji cells, as determined by flow cytometry and anti-CD22 staining. Quantification was performed with Quantibrite™ PE beads and data analysis assumed monovalent antibody binding. Statistical analysis was performed using a repeated measure One-Way ANOVA with a Tukey post hoc test (*$p < 0.05$, **$p < 0.01$, ***$p < 0.001$, ****$p < 0.0001$). The statistical analysis for the cytokine concentration was performed using log-transformed values.

Fig. S6

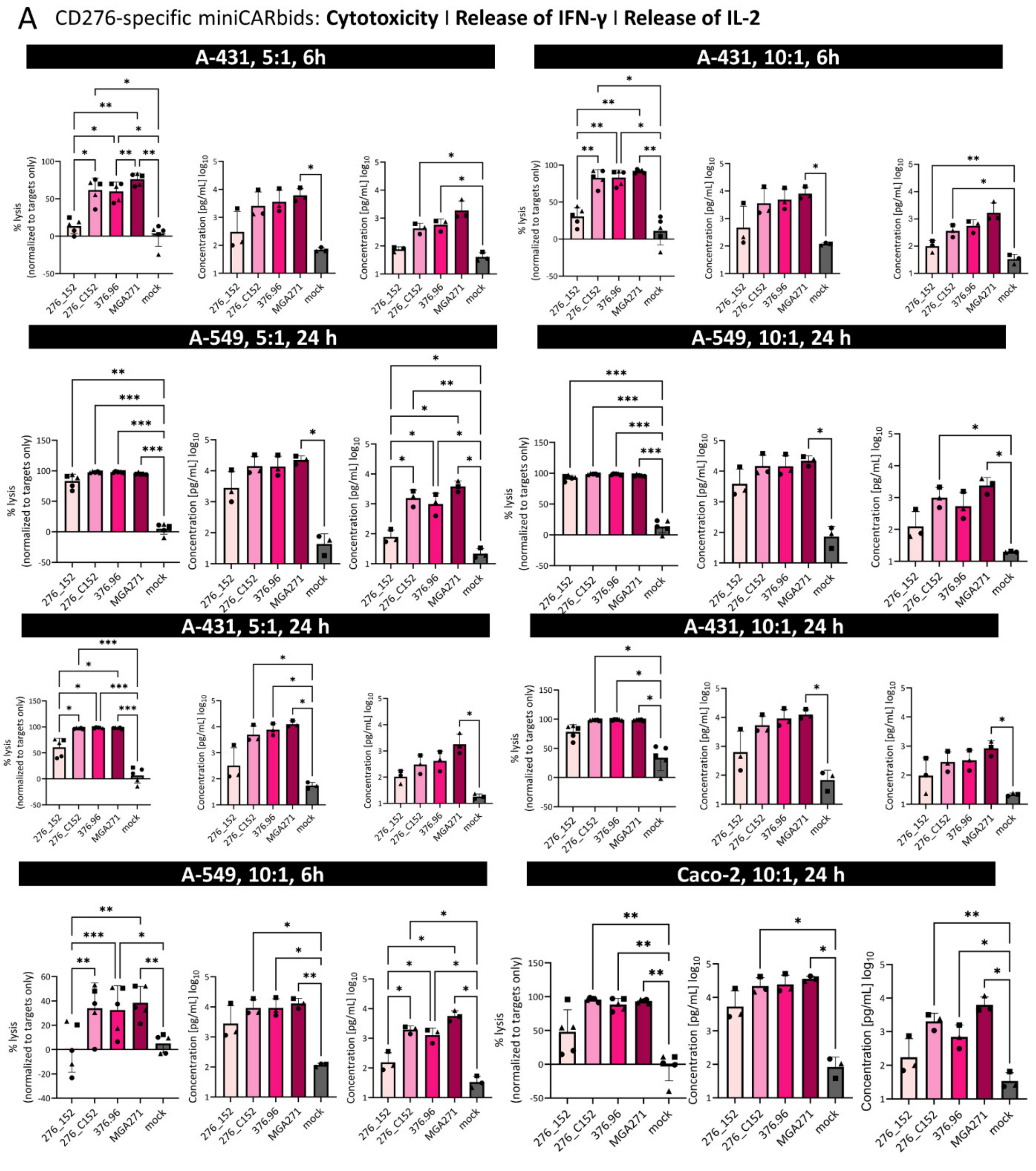


**Suppl. Fig. 6: Functional analysis of CD276-specific miniCARbid-CAR Ts.** Cytotoxicity of CD276-specific CAR T cells and mock T cells (no CAR) against A-431, Caco-2 or A-549 cells (E:T 5:1 or 10:1, 6 or 24 h, average ± SD, n=5, biological replicates). IFN-γ and IL-2 in the respective supernatants were analyzed via ELISA (average ± SD, n=3, biological replicates). Statistical analysis was performed using a repeated measure OneWay ANOVA with a Tukey post hoc test (*p < 0.05, **p < 0.01, ***p < 0.001, ****p < 0.0001). The statistical analysis for the cytokine concentration was performed using log-transformed values.

Fig. S7

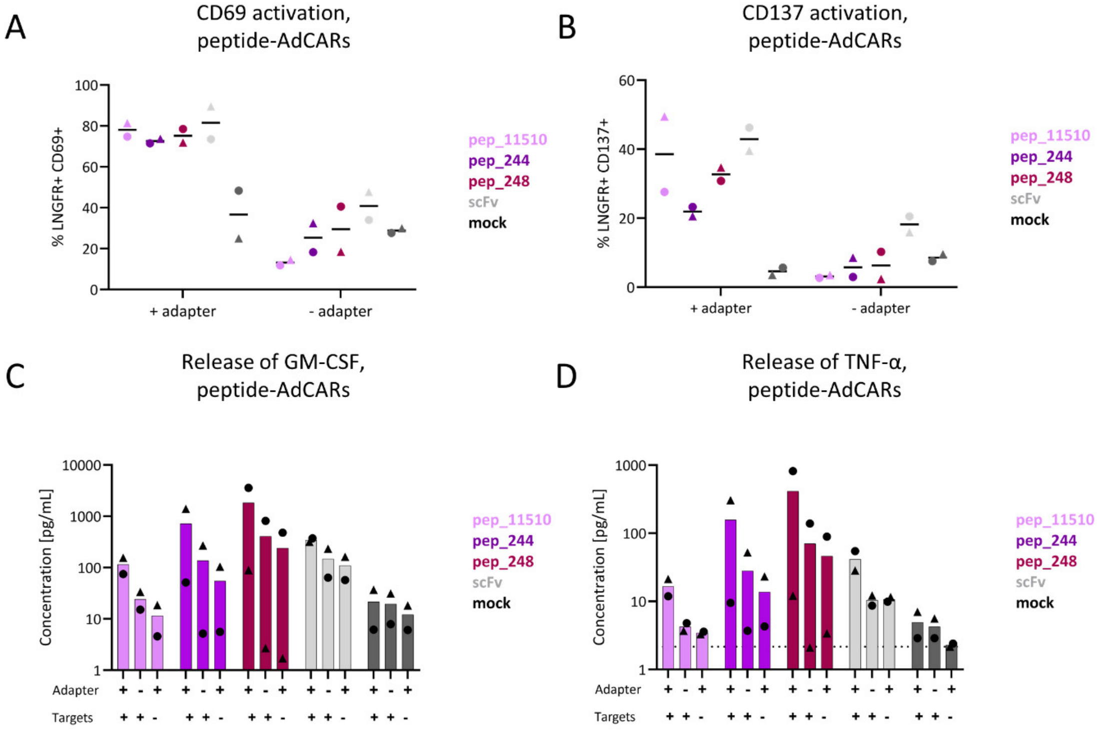


**Suppl. Fig. 7: Functional analysis of adapter CARs (AdCARs) based on peptide-specific miniCARbids.** (A) Analysis of CD69 expression on primary human T cells expressing peptide-specific AdCARs (three miniCARbid-CARs and one scFv-CAR as a benchmark). Untransduced (UTD) cells were measured as a control (n=2, biological replicates). Activation was measured with and without the addition of adapter protein. (B) Analysis of CD137 expression on primary human T cells expressing peptide-specific AdCARs. Untransduced (UTD) cells were measured as a control (n=2, biological replicates). Activation was measured with and without the addition of adapter protein. (C and D) Release of GM-CSF (C) and TNF-α (D) by peptide-specific AdCAR T cells as analyzed by MACSPlex Cytokine Kit (n=2, biological replicates). Release was tested under several conditions: +/+ (with adapter and target cells), +/- (with adapter and without target cells) and -/+ (without adapter and with target cells). The dotted line indicates the detection limit.

Fig. S8

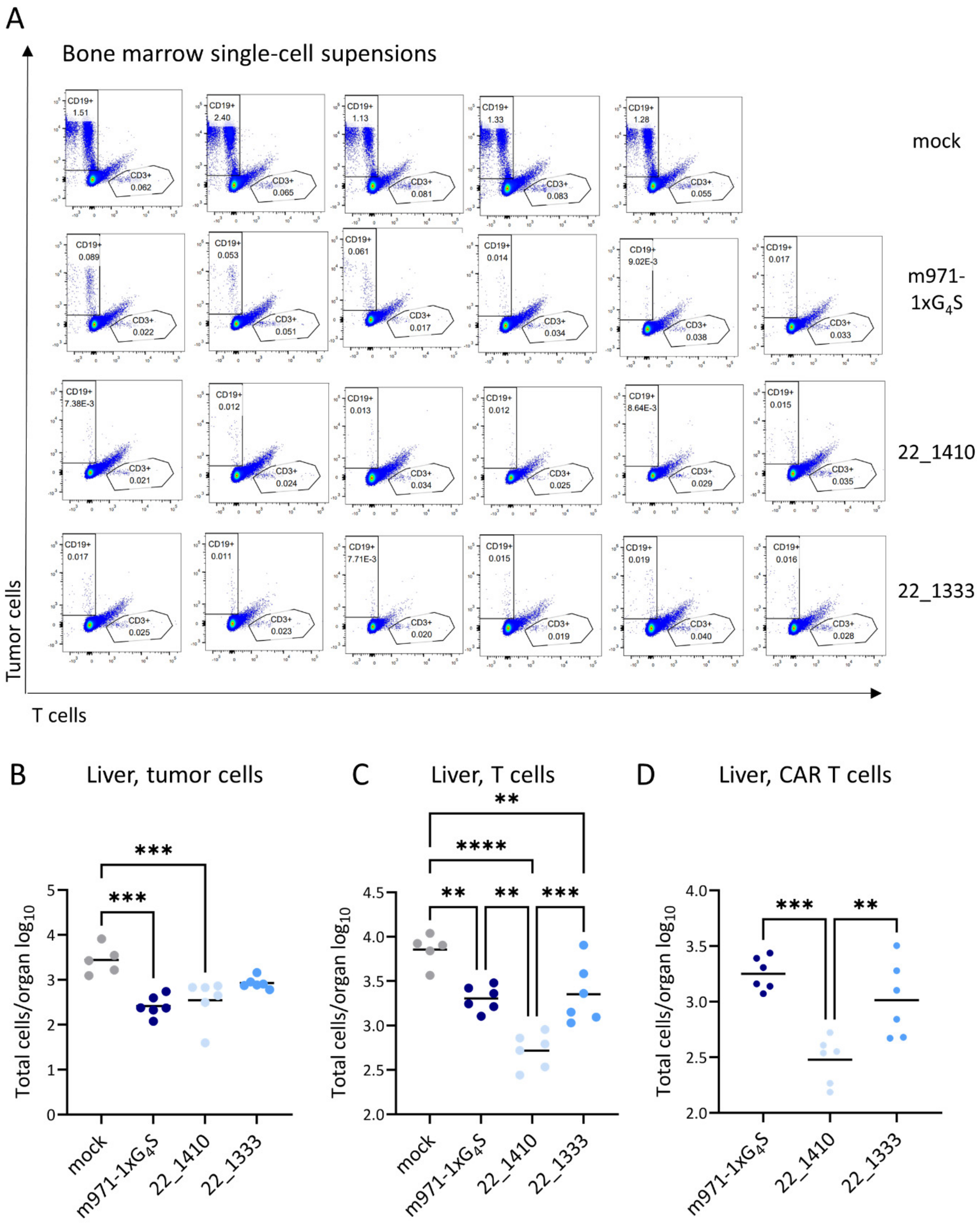


**Suppl. Fig. 8: Anti-tumor activity of CD22-directed miniCARbid-CAR Ts *in vivo*.** (A) Flow cytometric analysis of bone marrow single-cell supensions showing the percentage of tumor and CD3+ cells, respectively, for all mice infused with either mock T cells (no CAR), T cells expressing scFv-CAR m971-1x$G_4$S or T cells expressing miniCARbid-CARs 22_1410 or 22_1333. Each dot plot represents the flow cytometric staining of the bone marrow of one single mouse. (B) Analysis of the number of tumor cells (CD19+ GFP+) of five (mock T cells) or six animals (CAR expressing T cells) in the liver. Depicted is the log-transformed number of total cells in the liver. Each circle represents the cell number in one animal. (C and D) Analysis of the number of T cells (CD3+) (C) and CAR T cells (MAP+) (D) of five (mock T cells) or six animals (CAR expressing T cells) in the liver. Depicted is the log-transformed number of total cells in the liver. Each circle represents the cell number in one animal. Statistical analysis was performed using a One-Way ANOVA of log-transformed values with a Tukey post hoc test (**p < 0.01, ***p < 0.001, ****p < 0.0001).

### Fig. S9

#### A AA sequences of CD22 miniCARbids

```
5UMR       MAETLEFNDVYQEVKGSMNDGRLRLSRQGIIFKNSKTGKVDNIQAGELTEGIWRRVALGHGLKLLTKNGHVYKYDGFRESEFEKLSDFFKTHYRLELMEK
22_154     MAETLEFNDVYQEVKGSMNDGKLKLHRDVITFWNSKTGKYDWIWAGELTEGIWRRVALGHGLKLLTKNSHVYKYDGFRESEFEKLSDFFKTHYRLELMEK
22_1410    MAETLEFNDVYQEVKGSMNDGKLKLHRDVITFWNSKTGKYDWIQAGELTEGIWRRVALGHGLKLLTKNGHVYKYDGFRESEFEKLSDFFKTHYRLELMEK
22_163     MAETLEFNDVYQEVKGSMNDGKLKLHRDVITFWNSKTGKYDWIHAGELTEGIWRRVALGHGLKLLTKNGHVYKYDGFRESEFEKLSDFFKTHYRLELMEK
22_1611    MAETLEFNDVYQEVKGSMNDGKLKLHRDVITFWNSKTGKYDWIWAGELTEGIWRRVALGHGLKLLTKNGHVYKYDGFRESEFEKLSDFFKTHYRLELMEK
22_1324    MAETLEFNDVYQEVKGSMNDGKLHLMRDHIDFWNSKTGKYDWIQAGELTEGIWRRVALGHGLKLLTKNGHVYKYDGFRESEFEKLSDFFKTHYRLELMEK
22_164     MAETLVFNDVYQEVKGSMNDGKLHLMRDHIDFWNSKTGKYDWIWAGELTEGIWRRVALGHGLKLLTKNGHVYKYDGFRESEFEKLSDFFKTHYRLELMEK
22_1415    MAETLEFNDVYQEVKGSMNDGKLRLDREHIDFWNSKTGKYDWIWAGELTEGIWRRVALGHGLKLLTKNGHVYKYDGFRESEFEKLSDFFKTHYRLELMEK
22_1317    MAETLEFNDVYQEVKGSMNDGVLKLDRQHIDFWNSKTGKYDWIQAGELTEGIWRRVALGHGLKLLTKNGHVYKYDGFRESEFEKLSDFFKTHYRLELMEK
22_1316    MAETLEFNDVYQEVKGSMNDGVLQLDRQHIDFWNSKTGKYDWIYAGELTEGIWRRVALGHGLKLLTKNGHVYKYDGFRESEFEKLSDFFKTHYRLELMEK
22_1333    MAETLEFNDVYQEVKGSMNDGRLYLDRQHIDFYNSKTGKYDWIWAGELTEGIWRRVALGHGLKLLTKNGHVYKYDGFRESEFEKLSDFFKTHYRLELMEK
```

#### B AA sequences of CD276 miniCARbids

```
276_C146   MAETLEFNDVYQEVKGSMNDGQLHLMRSGIWFWNSKTGKLDYIYASELTEGIWRRVALGHGLKLLTKNGHVYRYDGFRESEFEKLSDFFKTHYRLELMEK
276_C152   MAETLKFNDVYQEVKGSMNDGQLYLQRSGIWFFNSKTGKLDIIYAGELTEGIWRRVALGHGLKLLTKNGHVYKYDGFRESEFEKLSDFFKTHYRLELMEK
5UMR       MAETLEFNDVYQEVKGSMNDGRLRLSRQGIIFKNSKTGKVDNIQAGELTEGIWRRVALGHGLKLLTKNGHVYKYDGFRESEFEKLSDFFKTHYRLELMEK
276_165    MAETLEFNDVYQEVKGSMNDGTLKLKRGIIVFWNSKTGKADMIIAGELTEGIWRRVALGHGLKLLTKNGHVYKYDGFRESEFEKLSDFFKTHYRLELMEK
276_158    MAETLEFNDVYQEVKGSMNDGTLLLYRGLIRFWNSKTGKYDFIIAGELTEGIWRRVALGHGLKLLTKNGHVYKYDGFRESEFEKLSDFFKTHYRLELMEK
276_152    MAETLEFNDVYQEVKGSMNDGQLKLYRGLIRFYNSKTGKYDFIIAGELTEGIWRRVALGHGLKLLTKNGHVYKYDGFRESEFEKLSDFFKTHYRLELMEK
276_1710   MAETLEFNDVYQEVKGSMNDGVLKLYRGLIRFYNSKTGKYDMIIAGELTEGIWRRVALGHGLKLLTKNGHVYKYDGFRESEFEKLSDFFKTHYRLELMEK

3SHU       SMKLVKFRKGDSV--GLRLAGGNDVGIFVAGVLEDSPAAKEGLEEGDQILRVNNVDFTNIIREEAVLFLLDLPKGEEVTILAQKKKDVYRRIVE
276_155    SMKLVKFRKGDQVWVSLYLAGGNDVGIFVAGVEEDSPAAKEGLEEGDQILRVNNVDFTNIIYHEAMYFLWDLPKGEEVTILAQKKKDVYRRIVE
276_173    SMKLVKFRKGDAVWLSLELAGGNDVGIFVAGVDEDSPAAKEGLEEGDQILRVNNVDFTNIIYGEAMYFLWDLPKGEEVTILAQKKKDVYRRIVE
```

#### C AA sequences of peptide miniCARbids

```
3SHU       SMKLVKFRKGDSVGLRLAGGNDVGIFVAGVLEDSPAAKEGLEEGDQILRVNNVDFTNIIREEAVLFLLDLPKGEEVTILAQKKKDVYRRIVE
pep_1056   SMKLVKFRKGDYVGLELAGGNDVGIFVAGVDEDSPAAKEGLEEGDQILRVNNVDFTNIIGWEALPFLLDLPKGEEVTILAQKKKDVYRRIVE
pep_1146   SMKLVKFRKGDYVGLELL-GNDVGIFVAGVDEDSPAAKEGLEEGDQILRVNNVDFTNIISMEALYFLLDLPKGEEVTILAQKKKDVYRRIVE
pep_1159   SMKLVKFRKGDYVGLDLA-GGDVGIFVAGVDEDSPAAEEGLEEGDQIMRVNNVDFTNIIHWEALDFLLDLPKGEEVTILAQKKKDVYRRIVE
pep_11510  SMKLVKFRKGDYVGLELA-GGDVGIFVAGVDEDSPAAKEGLEEGDQILRVNNVDFTNIISWEALYFLLDLPKGEEVTILAQKKKDVYRRIVE
pep_2215   SMKLVKFRKGDYVGLELA-GGDVGIFVAGVDEDSPAAKEGLEEGDQIMRVNNVDFTNIIHWEALYFLLDLPKGEEVTILAQKKKDVYRRIVE
pep_244    SMKLVKFRKGDYVGLELL-GNDVGIFVAGVDEDSPAAEEGLEEGDQIMRVNNVDFTNIIHWEALYFLLDLPKGEEVTILAQKKKDVYRRIVE
pep_248    SMKLVKFRKGDYVGLELL-GNDVGIFVAGVDEDSPAAKEGLEEGDQIMRVNNVDFTNIIHWEALYFLLDLPKGEEVTIQAQKKKDVYRRIVE
pep_245    GMMLVKFRKGDYVGLELL-GNDVGIFVAGVDEDSPAAKEGLEEGDQIMRVNNVDFTNIIHWEALYFLLDLPKGEEVTILAQKKKDVYRRIVE
```

**Suppl. Fig. 9: Amino acid (AA) sequences of miniCARbids.** Residues which are part of the mutated binding surface are displayed as bold letters in the respective WT sequences (5UMR or 3SHU, respectively). WT residues are highlighted in yellow, while the most frequent mutation on each position is highlighted in pink. Alignments of AA sequences of CD22-specific miniCARbids (A), CD276-specific miniCARbids (B) and peptide-specific miniCARbids (C) together with their respective parental proteins 3SHU and 5UMR using Clustal Omega.

**Suppl. Table 1**

| | |
|---|---|
| 5UMR_c_NNK1 | 5'-GAAGGGCAGCATGAACGACGGC**NNK**CTG**NNK**CTC**NNK**AGACAG**NNK**ATC**NNK**TTC**NNK**AACAGCAAGACCGGCAAG**NNK**GAC**NNK**ATCCAGGCCGGCGAACTGACCGAAGG-3' |
| 5UMR_cp_NNK1 | 5'-GAAGGGCAGCATGAACGACGGC**NNK**CTG**NNK**CTC**NNK**AGA**NNKNNK**ATC**NNK**TTC**NNK**AACAGCAAGACCGGCAAG**NNK**GAC**NNK**ATC**NNK**GCCGGCGAACTGACCGAAGG-3' |
| 5UMR_cy_NNK1 | 5'-GAAGGGCAGCATGAACGACGGC**NNK**CTG**NNK**CTC**NNK**AGACAG**NNK**ATC**NNK**TTC**NNK**AAC**NNK**AAGACC**NNK**AAG**NNK**GAC**NNK**ATCCAGGCCGGCGAACTGACCGAAGG -3' |
| 5UMR_PCR1_rev | 5'-CTTTTCCATCAGTTCCAGCCGGTAGTGG-3' |
| 5UMR_PCR2_fwd | 5'-GGAGGCGGTAGCGGAGGCGGAGGGTCGGCTAGCATGGCCGAGACACTGGAATTCAACGACGTGTACCAAGAAGTGAAGGGCAGCATGAACGACGGC-3' |
| 5UMR_PCR2_rev | (5'-GTCCTCTTCAGAAATAAGCTTTTGTTCGGATCCCTTTTCCATCAGTTCCAGCCGGTAGTGG-3' |
| 5UMR_cb_NNK2 | 5'-GGAGGCGGTAGCGGAGGCGGAGGGTCGGCTAGCATGGCCGAGACACTG**NNK**TTC**NNK**GACGTGTACCAAGAAGTGAAGGGCAGCATGAACGACGGC-3' |
| 3SHU_1_fwd | 5'-CTGGTCAAGTTCCGGAAGGGCGAT**NNK**GTG**NNK**CTG**NNK**CTG**NNK**GGCGGAAACGATGTGGGCATTTTTGTGGCTGGCGTGCTGGAAGATAGCCCTGCCGCCAAAGAAGG-3' |
| 3SHU_2_fwd | 5'-CTGGTCAAGTTCCGGAAGGGCGATAGCGTGGGACTG**NNK**CTG**NNK**GGCGGAAACGATGTGGGCATTTTTGTG**NNK**GGCGTG**NNK**GAAGATAGCCCTGCCGCCAAAGAAGG-3' |
| 3SHU_3_fwd | 5'-CTGGTCAAGTTCCGGAAGGGCGAT**NNK**GTG**NNK**CTG**NNK**CTGGCTGGCGGAAACGATGTGGGCATTTTTGTGGCTGGCGTG**NNK**GAAGATAGCCCTGCCGCCAAAGAAGG-3' |

| | |
|---|---|
| 3SHU_4_fwd | 5’-CTGGTCAAGTTCCGGAAGGGCGAT**NNK**GTG**NNK**CTG**NNK**CTGGCTGGCGGAAACGATGTGGGCATTTTTGTG**NNK**GGCGTG**NNK**GAAGATAGCCCTGCCGCCAAAGAAGG-3’ |
| 3SHU_1_rev | 5’-GTCACTTCCTCGCCTTTAGGCAGGTC**MNN**CAGGAA**MNNMNN**GGCCTC**MNNMNN**GATGATGTTGGTGAAGTCCACGTTGTTCACTC-3’ |
| 3SHU_4_rev | 5’-GTCACTTCCTCGCCTTTAGGCAGGTC**MNN**CAGGAA**MNNMNN**GGCCTCTTCTCTGATGATGTTGGTGAAGTCCACGTTGTTCACTC-3’ |
| 3SHU_PCR2_fwd | 5’-GGAGGCGGTAGCGGAGGCGGAGGGTCGGCTAGCAGCATGAAGCTGGTCAAGTTCCGGAAGGGCGAT-3’ |
| 3SHU_PCR2_rev | 5’-GTCCTCTTCAGAAATAAGCTTTTGTTCGGATCCTTCCACGATCCGCCGGTAGACGTCCTTCTTCTTCTGGGCCAGAATGGTCACTTCCTCGCCTTTAGGCAGGTC-3’ |
| 3SHU_lib3_PCR1_fwd | 5‘-CTGGTCAAGTTCCGGAAGGGCGAT **X01** GTG **X01** CTG **X01** CTGGCTGGCGGAAACGATGTGGGCATTTTTGTGGCTGGCGTG **X01** GAAGATAGCCCTGCCGCCAAAGAAGG-3‘ |
| 3SHU_lib3_PCR1_rev | 5‘-GTCACTTCCTCGCCTTTAGGCAGGTC **Z01** CAGGAA **Z01** **Z01** GGCCTC **Z01** **Z01** GATGATGTTGGTGAAGTCCACGTTGTTCACTC-3‘ |
| 5UMR_cp_PCR1_fwd | 5‘-GAAGGGCAGCATGAACGACGGC **X01** CTG **X01** CTC **X01** AGA **X02** **X01** ATC **X01** TTC **X01** AACAGCAAGACCGGCAAG **X01** GAC **X01** ATC **X02** GCCGGCGAACTGACCGAAGG-3‘ |
| SUMO-peptide fusion protein | DSEVNQEAKPEVKPEVKPETHINLKVSDGSSEIFFKIKKTTPLRRLMEAFAKRQGKEMDSLRFLYDGIRIQADQAPEDLDMEDNDIIEAHREQIGG<u>GGGGSYVVERWRHRP</u> |
